# Cell-free protein synthesis as a method to rapidly screen machine learning-directed protease variants

**DOI:** 10.1101/2025.01.24.634768

**Authors:** Ella Lucille Thornton, Jeremy T. Boyle, Nadanai Laohakunakorn, Lynne Regan

## Abstract

Machine learning (ML) tools have revolutionised protein structure prediction, engineering and design, but the best ML tool is only as good as the training data it learns from. To obtain high quality structural or functional data, protein purification is typically required, which is both time and resource consuming – especially at the scale required to train ML tools. Here, we showcase cell-free protein synthesis (CFPS) as a straightforward and fast tool for screening and scoring the activity of protein variants for ML workflows. We demonstrate the utility of the system by improving the kinetic qualities of a protease. By rapidly screening just 48 random variants to initially sample the fitness landscape, followed by 32 more targeted variants, we identified several protease variants with improved kinetic properties.

## Introduction

The success of machine learning (ML) methods depends on the quality of the data on which an algorithm is trained. The vast amount of high-quality structural data in the protein database (PDB) underlies the remarkable success of ML program AlphaFold in predicting protein structure from sequence [1]. To successfully apply ML methods to other areas of protein science, such as the design of function, high quality data is vital [2], [3].

Protein design space can be imagined as a landscape, populated with many different protein sequences each associated with a different ‘fitness’. The user-defined fitness could be any desirable characteristic such as binding affinity, catalysis rate, yield, or solubility (Figure 1) [4]. Identifying the protein with the highest fitness within this landscape can be facilitated by using a ML approach to sample and learn from high quality data from different sequences [5]. Employing statistical methods embedded in ML workflows helps the user to avoid common ‘traps’ in the navigation of a protein fitness landscape, such as getting stuck at local fitness optima: where a variant is better than original, but not the absolute best [6], [7]. It enables more efficient sampling of the wider fitness landscape to improve the likelihood of identifying the absolute best variant possible.

**Figure 1.**
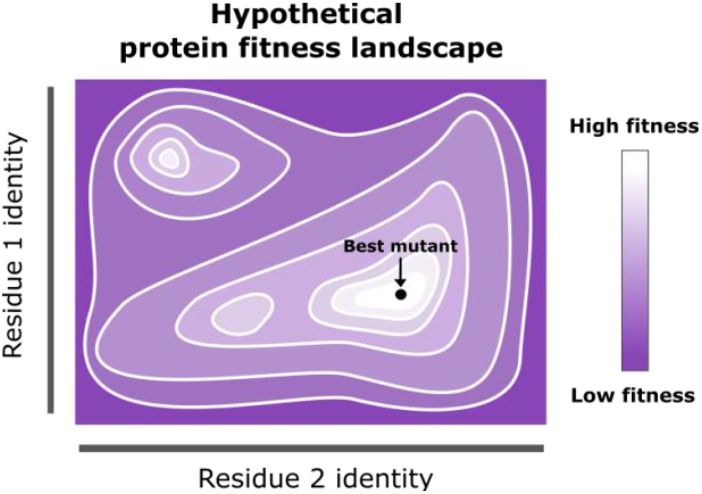
Diagram representing a hypothetical protein fitness landscape, where sequence identity is mapped to fitness. In this hypothetical scenario, two residues (residue 1 and 2) are selected for mutagenesis in the protein and can each be mutated to any other amino acid, represented on the ‘x and y axes’ of this space. Areas shaded with lighter colours represent protein variants with higher ‘fitness’, where fitness can refer to any protein functionality, for example: affinity, catalysis rate, yield or solubility. Labelled with an arrow is the best hypothetical variant.

Collecting functionality data for protein variants to explore the protein fitness landscape is often achieved by the time-consuming process of *in vivo* expression, purification and characterisation. But as ML-driven protein mutagenesis projects typically require 10-100’s of variants for the model to be effectively trained, a higher-throughput approach is required [8].

Many different experimental approaches have been previously employed to find Tobacco Etch Virus (TEV) protease variants with improved rates of catalysis. Sanchez & Ting use a directed evolution approach coupled to a high throughput yeast FACS workflow to identify uTEV, which has an improved catalytic rate [9]. Another system [10] uses yeast endoplasmic sequestration screening to test variants of TEV produced by error-prone PCR, finding eTEV, which has an improved rate of catalysis derived from an increase in turnover rate. Sumida *et al*. express and purify *de novo* TEV designs at a small scale, identifying several designs with improved stability and functionality compared to the starting TEV [11]. These examples demonstrate different approaches to protease design and screening, and all methodologies are successful in identifying new functional proteases, each with their own distinct advantages. Here, we detail a new protease-screening workflow, combining the use of ML and cell-free protein synthesis (CFPS).

CFPS is a powerful tool for protein production and functionality screening [12], [13], [14]. By removing the machinery from the cell required for transcription and translation, it can be uncoupled from cellular survival and the scope for applications is significantly widened [15], [16]. For example, it enables the production of proteins incompatible with cell survival, or the use of substrates impermeable to the cell wall [17]. Because CFPS is an open system, the reaction conditions are more easily programmable compared to living cells: ionic strength, pH and redox potential can be controlled for production of specific proteins [18].

We demonstrate CFPS’s role in the protein ML workflow by rapidly assessing 100’s of protease variants for functionality by combining CFPS with an assay for protease activity. Providing this sequence-fitness data to an ML algorithm [19] to find optimal variants, we identify several active variants of our test protease and improve the overall fitness of the protease by 4-fold. The throughput of the system is demonstrated by the ability to produce and screen an initial round of 48 variants within 6 hours.

## Results

Here we present the results of combining CFPS with ML to identity variants of a protease with enhanced activity. The protein of interest for this campaign is Con1 [20], a designed protease created for the removal of tags from proteins after purification. Con1 exhibits high solubility and desirable cleavage specificity, in comparison to TEV. Our goal was to identify variants of the Con1 protease that display enhanced fitness, defined in this instance as the increase in initial rate of substrate cleavage at a single fixed substrate concentration.

We first used AlphaFold3 [21] to model the Con1 protease with substrate bound (Figure 2). We then used Robetta alanine scan [22] to identify residues predicted to contribute significantly to substrate binding. Six were identified: H167, L169, F172, L217, Q218 and E219, which are predicted to fall on opposite sides of the substrate. We therefore split our mutagenesis scheme to target each of these regions independently, with Region A consisting of H167, L169, F172, and Region B consisting of L217, Q218, E219 (Figure 2).

**Figure 2.**
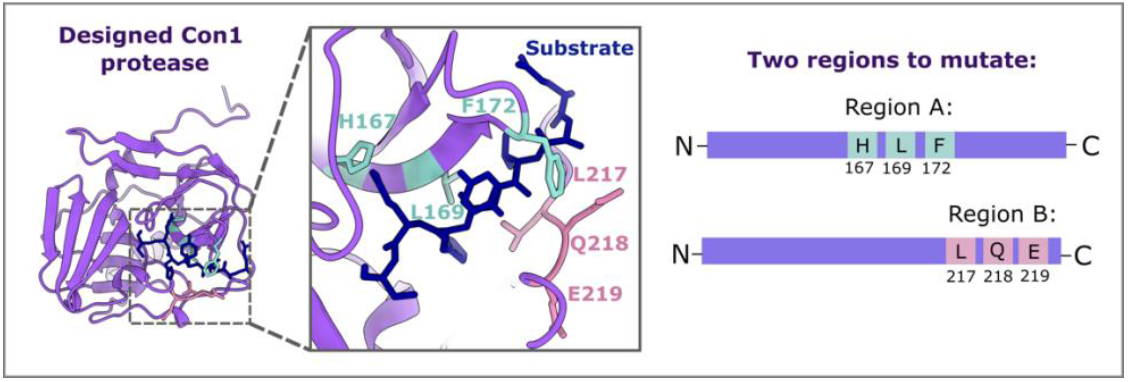
Residues to mutate within the Con1 protease were identified by computational alanine scanning. Six residues were selected and split into two regions to be mutated simultaneously: region A (blue) or B (pink).

We created 48 protein variants per Region (A or B), where each of the three residues were randomly mutated to any other amino acid (Figure 3). DNA templates encoding each of these variants were combined with cell-free reagents (lysate, amino acids, additives), and the proteases were synthesised *in vitro*.

**Figure 3.**
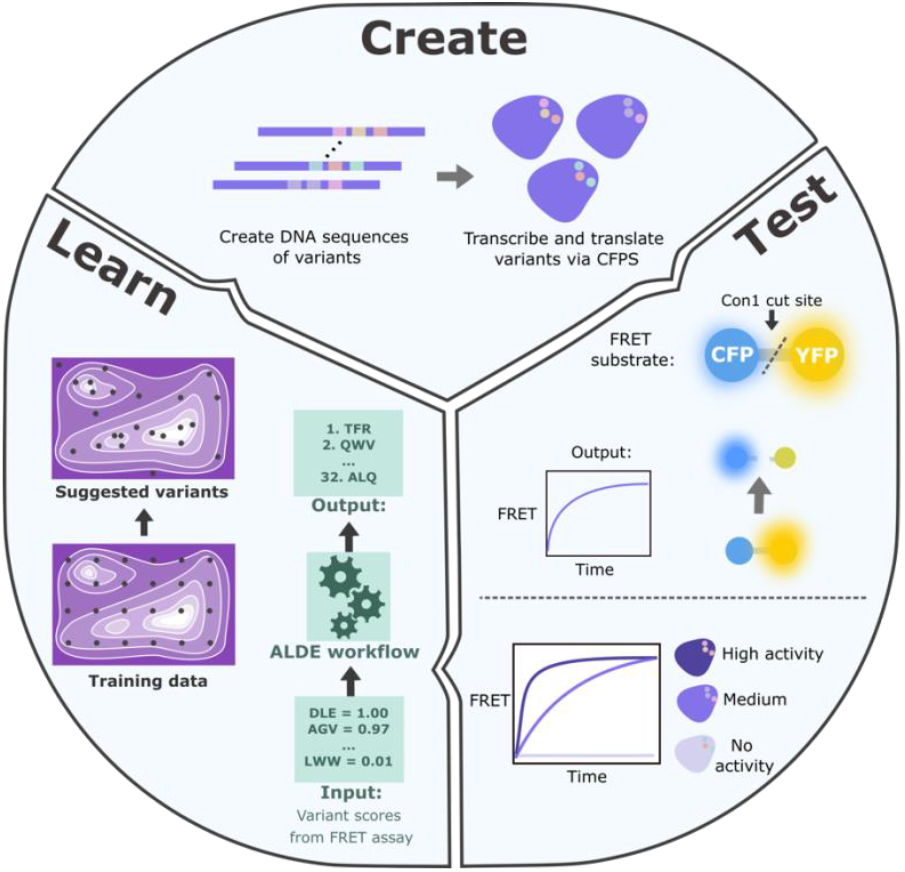
Overview of workflow. The process is split into three phases: (1) Create, (2) Test, (3) Learn. First, 48 random variants were created per region within the protease (A or B, as specified in Figure 2). Linear DNA encoding for these variants was combined with CFPS reagents and incubated at 37 °C to produce the protease variants *in vitro*. Next, protease variants were tested for activity by addition of FRET substrate and monitoring an increase in FRET over time. The catalytic ‘speed’ of each variant was recorded and translated to a fitness score normalised to WT. This sequence-fitness data was provided to the ALDE ML workflow [19] as training data. The workflow then provides suggested variants to best explore the fitness landscape. Linear DNA encoding 32 of these suggested variants was then combined with CFPS reagents for in vitro expression, and the cycle of Create-Test-Learn continues.

The activity of each Con1 protease variant was assessed using a FRET-based assay. The substrate for this assay is a purified fusion protein in which CFP (donor) and YFP (acceptor) are linked by the specific Con1 protease substrate cut site. By monitoring fluorescence of both the donor and acceptor protein, a FRET ratio can be calculated and tracked over time, providing a measure of the substrate cleavage rate for each variant (Figure 3).

### Sparse sampling of the fitness landscape provides an insight to crucial residues required for function

Screening of the 48 variants for region A and region B revealed striking differences between the tolerance of each region to mutation. Figure 4 shows the variant sequences tested and their activity relative to the original Con1 protease.

**Figure 4.**
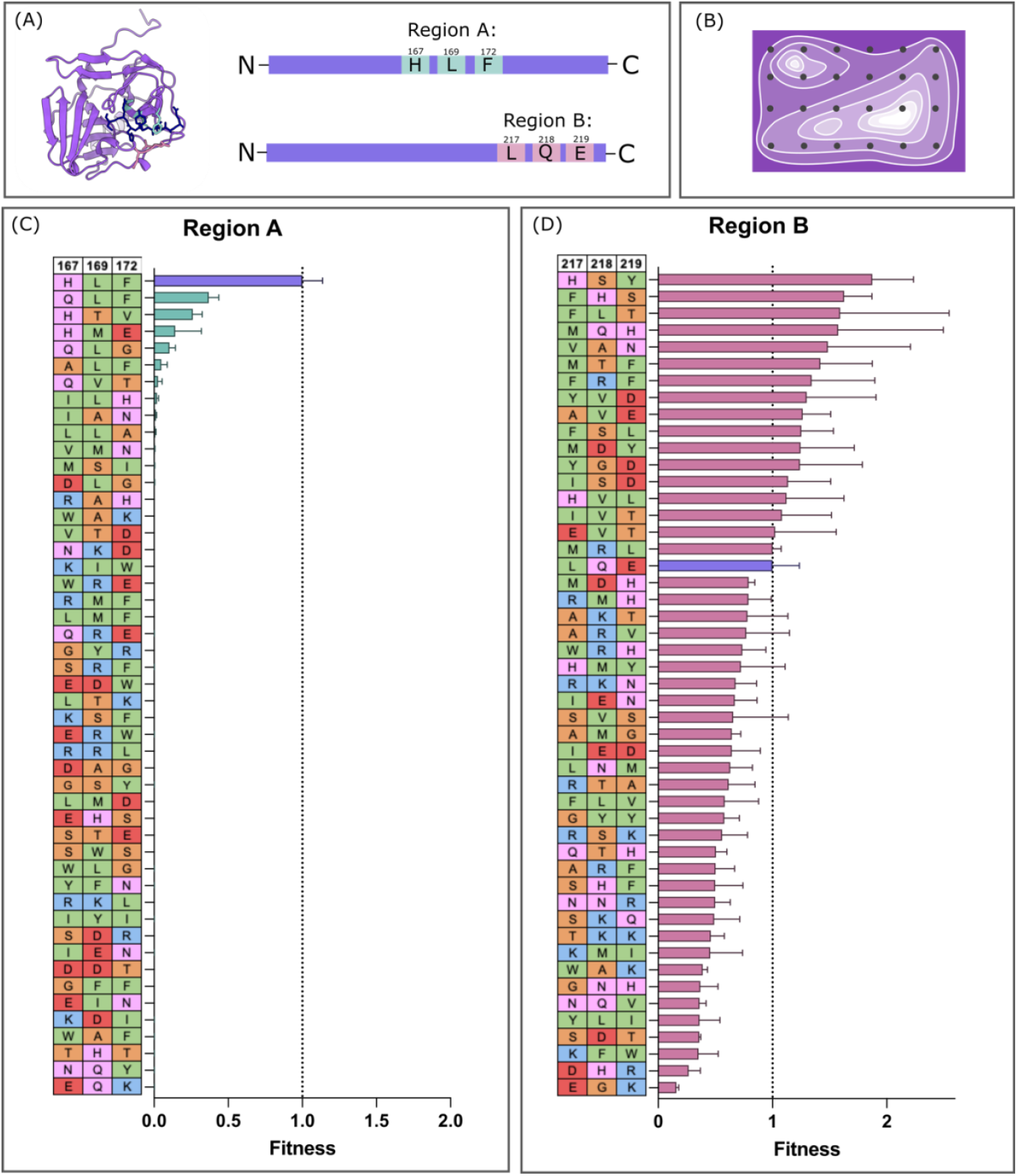
Sparse sampling of mutational landscape for Con1 protease. (A) Con1 protease AlphaFold model and cartoon schematic of the protease, with highlighted residues to be mutated. Region A includes 167H, 169L, 172F and region B includes 217L, 218Q, 219E. The two regions were mutated and tested independently. (B) 48 random variants were tested for both region A and region B. In the context of the protein fitness landscape, as introduced in Figure 1, this could be visualised as sparse sampling of the total available variants in the space. (C, D) Based on the speed of substrate cleavage, variants were scored, and values normalised to the original Con1 (shown in purple) to give a final fitness score for each. The identity of each variant at the residues to be mutated is indicated in the table, with amino acids coloured based on type: green = hydrophobic, pink = polar, red = negatively charged, orange = small non-polar, blue = positively charged. Error bars represent S.D. from mean with 3 replicates.

For region A, mutations are not well tolerated as seen by the overwhelming number of inactive variants (Figure 4C). Only a few out of the 48 tested show any activity, and these variants are very similar to starting identity, for example the best variant, A10, is QLF, which is very similar to the original sequence in these positions of HLF. In region A, none of the tested variants were more active than the original Con1.

For region B variants, we observed a much greater range of activity, with all variants exhibiting some activity against the substrate, and several exhibiting higher activities than that of the starting Con1 protease against the substrate. The mutations in regions A and B and their differing range of activities provide two contrasting sets of data to train the ML workflow on.

### ML suggested variants improve mean and maximum fitness

The sequence and corresponding fitness score for each variant served as training data for the ‘Active Learning-assisted Directed Evolution’ (ALDE) ML workflow [19]. Amino acid identity was recorded as a one-hot encoding, with no indication of the amino acid’s chemical nature. Each variant’s fitness score was specified by the initial rate of substrate cleavage.

Based on the training data, ALDE returned a list of variants to be tested next. The recommended variants navigate the protein fitness landscape by balancing exploitation of areas likely to contain high fitness variants with exploration of areas with uncertain predicted fitness. The normalised fitness scores of these variants alongside their identity is shown in Figure 5B and 5C.

**Figure 5.**
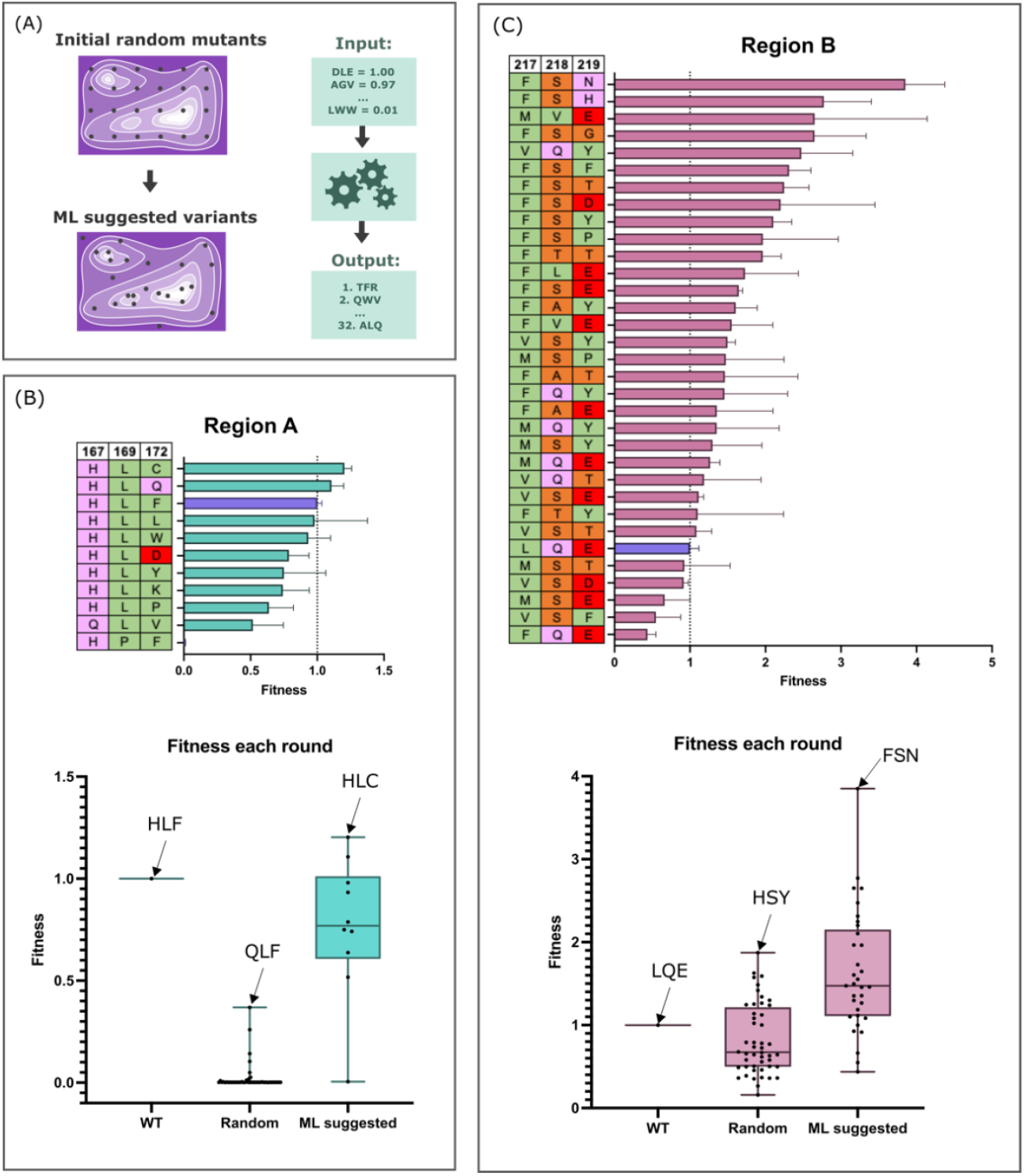
Suggested variants from ML workflow improve fitness of protease successfully. (A) Training data as shown in Figure 4 was input to the ML workflow, which then predicted the best next variants to test in the fitness landscape. The top ML predicted variants were made via CFPS and screened via FRET substrate, with 10 tested for region A (B), and 32 for region B (C), with fitness shown in ranked order for each mutant with corresponding identity. Error bars represent S.D. from mean with 3 replicates. Overall improvement of variant fitness is shown in each boxplot, where both the mean fitness and maximum fitness improves for both regions. The average fitness for each variant is shown as a single data point, with top variants at each step annotated with their identity.

Because random variants in region A were mostly non-functional, we hypothesised these residues were essential for protease function and intolerant to mutation. Since the likelihood of finding a fast variant seemed low, we only selected 10 ML-predicted variants to synthesise and screen. As shown in Figure 5B, the overall fitness was much improved, with both the mean fitness and maximum fitness improving from the random 48 variants. However, the top variants display fitness scores similar to the original Con1, and looking at the original identity (HLF) compared to the top variant in round 1 (QLF) with round 2 (HLL), there is a strong preference for original-like residues at these positions.

Region B variants were much more fruitful in terms of improved activity (Figure 5C). The top 32 predicted variants were synthesised and screened, with many faster than original Con1. Both the mean and maximum fitness improved compared to the random variants. The best performing variant (TB27) displayed fitness 4-fold higher than the original Con1 protease.

### Protease activity in CFPS reflects activity from purified and normalised proteases

To fairly compare the activity of different enzyme variants, they should be assayed at the same concentration. Because each CFPS reaction was provided with the same concentration of variant DNA, and the variants only differed by a maximum of three amino acids, we did not expect significant differences in expression levels. Nevertheless, it was still important to confirm that observed changes in protease activity were not due to differences in protease concentration.

We therefore selected four protease variants with varying levels of activity measured in CFPS screening (Figure 6a). These variants were expressed in *E. coli*, purified, normalised to the same concentration, and assayed using the FRET substrate (Figure 6B). As shown in Figure 6, comparable trends of activity are observed when the variants are screened either directly in CFPS, or as purified proteins at equal concentrations, therefore indicating that trends in variant activity in CFPS are reflecting genuine differences in protease activity.

**Figure 6.**
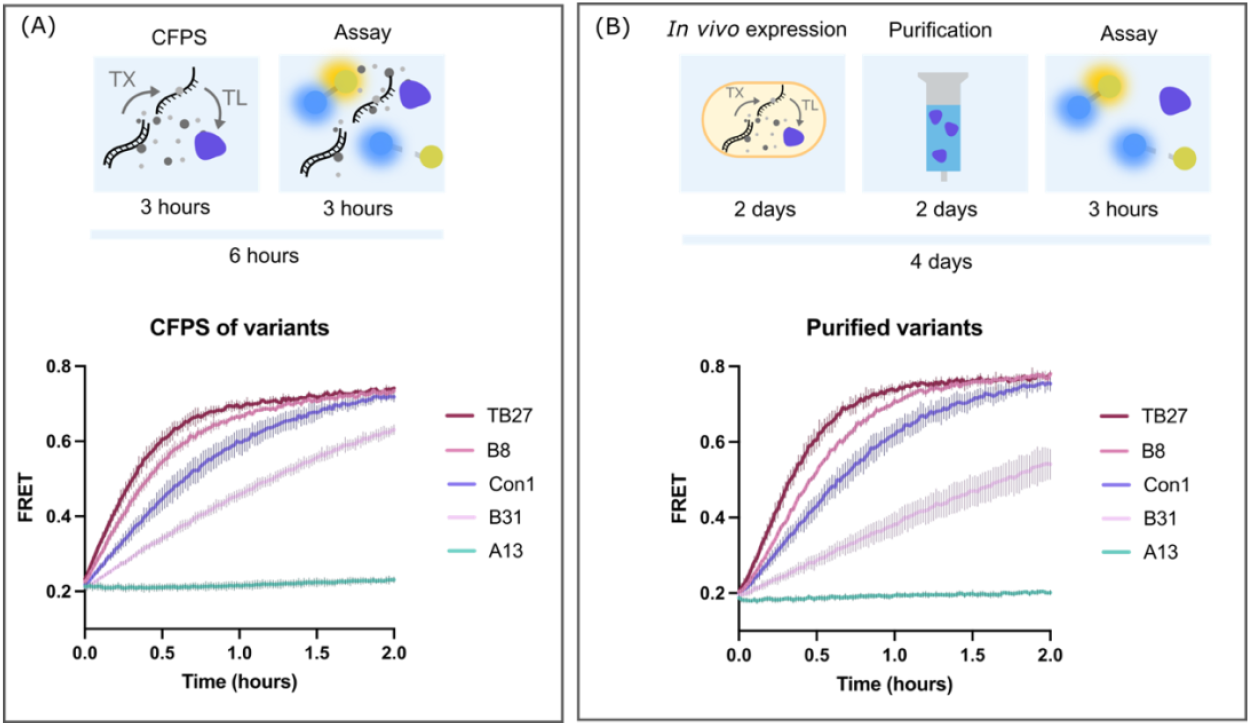
Comparison of protease activity assayed in CFPS with purified protein. (A) Four protease variants with varying apparent activity were selected from the CFPS screen and tested alongside the original Con1 protease. TB27 is the best performing variant from the ML suggested batch for region B variants, with a sequence identity of FSN at positions 217, 218, 219. B8 is the best performing variant from the initial random variants for region B, with identity HSY at 217, 218, 219. B31 is low performing variant from the initial random variants for region B, with identity TKK at 217, 218, 219. A13 is an inactive variant from the initial random variants for region A, with an identity V, M, N at 167, 169, 172. Shown here is the FRET output over time for each variant after production in cell-free. The time taken to obtain this data was 6 hours total. (B) The same four variants were transformed into BL21 *E. coli* cells and expressed, purified, quantified, normalised to 0.1 μM, and assayed. This took one working week. The output FRET over time is shown, and as in (A), the variants show similar trends of activity. Shown in both line plots is the mean signal from n=3, with S.D. shown as error bars.

## Discussion

Here, we present CFPS as a straightforward, fast, accessible tool for screening enzyme variants. Applied in microplate format as a medium-throughput technique (∼10-100’s), it can be strategically combined with ML tools to effectively navigate the fitness landscape and improve desired activity. This powerful workflow is demonstrated through the rapid discovery of an enzyme variant 4x faster than the starting sequence.

A key benefit of this workflow is the rapid speed at which an advantageous variant can be discovered. ML enables the efficient identification of high fitness variants within a complex fitness landscape by assessing combinations of mutations simultaneously, rather than in a stepwise ‘greedy’ fashion [7]. By combining this powerful analysis method with a fast and efficient data collection approach: CFPS coupled with protease activity screen, the cycle of create – test – learn can rapidly traverse the fitness landscape with minimal lab contact time required. The experimental side of the workflow could be sped up even further, by use of a faster commercial CFPS system, or by use of robotics to automate the pipetting [23], [24]. However, demonstrated here is an approach accessible to most labs to set up in-house for a relatively low cost, to find better variants of a protein of interest.

Perhaps the biggest obstacle to overcome in the process of beginning a protein design campaign is identifying an assay that can score your protein function accurately, and to this point we emphasise the unique benefits of using CFPS compared to cell-based assays: the inherently open system of CFPS allows for addition of a standard concentration of substrate that may be incompatible with living cells. Purification of the product from CFPS for further screening is also possible, by chromatography, or by surface capture [25], [26].

We show that activity observed directly from CFPS is comparable to activity observed with purified proteins. However, not every design campaign may be this straightforward, and normalising activity to variant expression levels is a desirable additional tool to this workflow. We would suggest the adaptation of one of the recently published protein tags for protein quantification in CFPS [27], [28], [29], [30] as an addition to the C-terminus of the designed protein to improve accuracy of activity scoring.

Enzyme design is not straightforward because there are many different features that contribute to enzymatic activity, not all of which have been identified for many classes of enzyme [31]. In our design campaign, we note that an exploratory approach to variant creation worked well, revealing dramatic differences between the tolerance of region A and region B for mutagenesis. These differences were not predictable at the computational analysis stage of variant design. Our work illustrates how such screening can provide information beyond that which can be obtained from analysis of sequence conservation. Evolution acts to identify the best sequence for a particular function in the context of the living organism. When a single enzyme is taken out of that context, the optimal sequence for the desired function is not necessarily the same. For example, position 167 of region A is conserved between similar proteases [20], but position 169 and 172 are not. Nevertheless, positions 167, 169 and 172 all display the same intolerance to mutation.

We demonstrate through CFPS paired with ML a 4-fold improvement of kinetic properties to our test protease. In total, 138 variants were rapidly screened in CFPS, each protein variant taking only 6 hours total to produce and screen, with 10’s screened simultaneously in the same plate. This work outlines an accessible and fast approach to find higher fitness variants of your protein of interest, and we envision this workflow can be adapted to many diverse applications of protein design.

## Methods

### General *E. coli* methods

Standard overnight cell growth: *E. coli* cells were picked from a single colony on an LB agar plate with the appropriate antibiotic into 5 mL LB with the appropriate antibiotic and grown overnight at 37 °C with shaking. For DNA preparation, TOP10 cells were used (F-*mcrA* Δ(*mrr*-*hsd*RMS-*mcr*BC) φ80*lacZ*ΔM15 Δ*lacX*74 *nup*G *recA*1 *araD*139 Δ(*ara-leu*)7697 *gal*E15 *gal*K16 *rps*L(StrR) *end*A1 λ-). For protein expression, BL21 Gold(DE3) cells were used (F-*omp*T *hsdSB*(*rB*–*mB*–) *dcm* (TetR) *gal* λ(DE3) *endA* The). For CFPS lysate production, Rosetta-gami2 cells were used (Δ(ara-leu)7697 ΔlacX74 ΔphoA PvuII phoR araD139 ahpC galE galK rpsL F′[lac+ lacIq pro] gor522::Tn10 trxB pRARE2 (Cam^R^, Str^R^, Tet^R^). Plasmids were transformed into competent *E. coli* cells following standard protocols [32].

DNA purification and quantification: Plasmids were purified from *E. coli* following protocols described by the manufacturer using the QIAprep® Spin Miniprep Kit (Qiagen). Linear DNA was purified from PCR mixture or agarose gels using the Promega Wizard® SV Gel and PCR Clean-up System following protocols described by the manufacturer. Purified DNA solutions were quantified by A260 and stored at -20 °C.

DNA sequencing: Plasmid sequences were verified by DNA sequencing performed either by DNA Sequencing & Services (www.dnaseq.co.uk) or Source Bioscience (www.sourcebioscience.com), using primers provided by the company.

### Design and cloning of variants

The identity of the random protease variants was decided by a python script which assembled 48 random variants with 3 identities per variant which could be any of the amino acids. Variants for each region were checked for complete amino acid representation (Figure S3). All designs were then created on Benchling and primers designed to change the codons for 3 amino acids specified in each variant.

All variants were synthesised by AQUA cloning [33], using the original Con1 protease sequence in pMAL plasmid as the template and mutating the DNA sequence via primer overhangs. All sequences and primers used are listed in the supplementary materials. Designed primer pairs for each Con1 variant were combined with the original Con1 plasmid for PCR by Phusion DNA polymerase, according to manufacturer’s instructions. After validation of successful DNA amplification by gel electrophoresis, DNA of each variant was purified from the PCR mixture by Zymo DNA cleanup kit. Pure DNA solution of each variant was then Dpn1 treated to remove any residual methylated plasmid. 0.5 μL DNA of each treated DNA (150 ng total), 0.5 μL FD buffer, 0.5 μL Dpn1, 3.5 μL H2O were combined in PCR tubes and incubated for 15 minutes at 37 °C, followed by 5 minutes at 80 °C to inactivate the enzyme. 2.5 μL of this treated mixture was combined with 25 μL of chemically competent TOP10 *E. coli* cells and incubated on ice for 15 minutes before heat shock at 42 °C for 45 seconds. After a brief incubation on ice, 200 μL LB was added to each and incubated at 37 °C for 20 minutes with shaking. As ampicillin was used as the selective antibiotic, short recovery of the cells was sufficient for growth. The total transformation mixture was plated on ampicillin (100 μg/mL final concentration) LB agar plates and incubated overnight at 37 °C. Generally, this protocol resulted in 10-40 colonies that displayed 95% successful identity (validated by sequencing).

### Cell-free protein synthesis

Lysate production: This protocol is adapted from Kwon & Jewett (2015). Typically, addition of 10 nM linear DNA to our homemade CFPS system yields 1 μM protease in a 5 μL reaction. 2xYTPG (16 g/L tryptone, 10 g/L yeast extract, 5 g/L NaCl, 7 g/L KH2PO4 3 g/L K2HPO4, 18 g/L glucose) was inoculated with 1/200 dilution of overnight cultures of Rosetta-gami2 *E. coli* cells. Cultures were grown for 2 hours at 37 °C with shaking, then induced with 0.4 mM IPTG and grown for a further 2 hours in the same conditions, before growth arrest by placing on ice. Cells were harvested by centrifugation at 10,000 xg for 10 minutes at 4 °C, the supernatant was discarded, and cell pellets were resuspended with 80 mL Buffer A (10 mM tris acetate (pH 8.2), 14 mM magnesium glutamate, 60 mM potassium glutamate) per 400 mL cells harvested. Cells were collected by centrifugation at 4,500 rpm for 10 minutes at 4 °C. The washing and cell harvesting process was repeated twice more, and cell pellets stored at -70 °C for future downstream processing. Cell pellets were resuspended with 1 mL buffer A per 1g wet cell mass and homogenised by vortexing. 1.5 mL aliquots were sonicated (Fisher120 W sonicator with probe for 0.5-15 mL) with pulses of 10s on, and 10s off, until a total energy output of 556 J was achieved, while incubated on ice. Lysate was clarified by centrifugation at 12,000 x g at 4 °C for 10 minutes. Supernatant was removed and centrifuged again to remove any residual insoluble material. Clarified supernatant was placed in a clean 1.5 mL tube and incubated at 37 °C for 1.5 hours with shaking (220 rpm) in a ‘run-off’ reaction, intended to allow the completion of translation of any mRNA associated with ribosomes. Samples were then centrifuged at 12,000 xg at 4 °C for 10 minutes. Supernatant was removed and aliquoted into 25 μL samples, which were stored at -70 °C until they were required for use.

Energy solution production: Energy solution was optimised for protease production via a Design of Experiments (DoE) approach. We used a central composite design with axial points to assess the best concentrations of Mg-glutamate, K-glutamate and PEG-8000. Further details about the DoE process and results are outlined in the supplementary information (Figure S1) Energy solution was assembled from stock solutions of all constituents, making a solution with a stock concentration 4x the working concentration of each component. In this 4x stock solution are the following components. Amino acid stock solution contained 50 mM each of the following amino acids: Alanine, Arginine, Asparagine, Aspartate, Cysteine, Glutamate, Glutamine, Glycine, Histidine, Isoleucine, Leucine, Lysine, Methionine, Phenylalanine, Proline, Serine, Threonine, Tryptophan, Valine. Tyrosine was prepared separately, in an acidic solution (pH ∼5.2) also at a final concentration of 50 mM. Stock batches of energy solution were prepared in volumes of 3 mL, with the following recipe: HEPES (pH 8) 200 mM, ATP 6 mM, GTP 6 mM, CTP 3.6 mM, UTP 3.6 mM, tRNA 0.8 mM, CoA 1.04 mM, NAD 1.32 mM, cAMP 3 mM, Folinic acid 0.27 mM, Spermidine 4 mM, 3-PGA 120 mM, Amino acids 6 mM, Tyrosine 3 mM, PEG-8000 8%, Mg-glutamate 31 mM, K-glutamate 880 mM, DTT 1 mM. Aliquots at 25 μl were then stored at -70 °C until required.

Preparation of DNA for cell-free reactions: DNA PCR product was used for CFPS reactions, with expression from a pTac promoter upstream of protein coding DNA sequence (full sequences available in supplementary table S1). DNA was purified by DNA Clean & Concentrator Kit (Zymo Research) following described protocols. Final DNA concentration was measured by absorbance at 260 nm on a Nanodrop (DeNovix DS-11) and DNA was stored at -20 °C until required.

Assembly of cell-free reaction: Cell free reactions were prepared with a final volume of 5 μL in wells of a low volume 384 microplate (Greiner, 788096). Master mix (MM) solutions were prepared for triplicates of each sample, with a final volume of 17 μL. Each MM contained: 6.13 μL lysate, 4.38 μL energy solution, 4.38 μL DNA (to give a final working concentration of 10 nM), 1.75 μL buffer A, 0.875 μL GamS nuclease inhibitor protein (NEB, P0774S). The MM was thoroughly mixed before 5 μL was deposited into each well using a multichannel pipette to dispense 12 sample MM at a time. The plate was sealed with an aluminium seal (Thermo, Z721557), which we found drastically reduces evaporation compared with plastic seals, and removes the need for wax. The plate was then incubated on a benchtop plate shaker at 37 °C with 1000 rpm shaking for 3.5 hours.

### FRET activity assay

Protease variants were assayed directly in CFPS reagents, or in purified form in buffer (100 mM Tris-HCl (pH 8.0), 25 mM NaCl, 5% glyercol) in a low volume 384 microplate (Greiner, 788096). 1 μL of purified FRET substrate was added to each 5 μL solution containing protease to give a final concentration of 5 μM. Substrate was added quickly to all wells using a multi-channel pipette to reduce the variability in reaction start time. The plate was sealed with an aluminium seal and quickly placed into the plate reader (POLARstar OMEGA) to measure fluorescence every 1 minute at (excitation/emission): (430/480), (430/520), both with gain set to 1000.

### Protein expression and purification

Cell growth and protein expression: Overnight cultures were diluted 100-fold into LB containing the appropriate antibiotic and grown at 37 °C with shaking until OD600 reached 0.6 – 0.8. Protein expression was induced by addition of Isopropyl β-d-1-thiogalactopyranoside (IPTG) to a final concentration of 1 mM and growth continued for a further 20 hours at 20 °C with shaking. Cells were collected by centrifugation at 6,000 x *g* for 10 minutes and pellets stored at -20 °C until needed.

Lysis and clarification: Cells were resuspended in lysis buffer containing cOmplete Protease Inhibitor Cocktail (Sigma-Aldrich) according to manufacturer’s instructions at a ratio of 1:50 volume of buffer to original cell culture volume. Resuspended cells were sonicated (Soniprep 150, MSE) on ice for 30 seconds, followed by a 30 second rest period. This sonication-rest cycle was repeated until cell lysis was achieved. Clarified cell lysates were prepared by centrifugation at 10,000 x *g* for 30 minutes at 4 °C.

Affinity purification: Proteins were purified via hexahistidine tag using Ni-NTA agarose (Qiagen) according to the manufacturer’s instructions. Purification was monitored using SDS-PAGE. Fractions containing protein at approximately 90% or greater purity, were pooled and dialysed against the desired storage buffer. Representative SDS-PAGE gel showing affinity purification of mutants is shown in figure S8.

Size exclusion chromatography (SEC): Further purification of proteins by size exclusion chromatography (SEC) with a Superdex-75 or Superdex-200 column was used when affinity chromatography did not provide sufficient purity. Purification by SEC is shown in Figure S9.

Protein quantification: Protein concentration was determined by measuring absorbance at 280 nm using the extinction coefficient of each protein calculated from the amino acid sequence using the tools available on Benchling [Biology Software].

Protein concentration: Buffer exchange and simultaneous concentration of protein solutions was performed using Amicon® Ultra Centrifugal Filters (Merck).

SDS-PAGE: Protein expression and purification were monitored using SDS-PAGE of samples alongside Precision Plus Protein Dual Xtra Prestained Protein Standards as molecular weight marker. Protein bands were visualised by staining using InstantBlue® Coomassie Protein Stain according to manufacturer’s instructions and were subsequently imaged using a Bio-Rad Gel Doc XR+ system.

### Data analysis and scoring of activity

FRET ratio was calculated by dividing blank (water) subtracted fluorescent signal at 480 nm by blank subtracted fluorescent signal at 520 nm, when solution was excited at 430 nm. This FRET ratio was plotted against time (GraphPad prism). In this project, we looked to increase the initial rate of substrate cleavage. To score the variants for this metric, the FRET data observed over time was truncated at FRET = 0.55, as this provided the best indication of initial rates of cleavage (Figure S4, S5, S6). The data was filtered using a python script. The filtered data was then plot in GraphPad, and a linear line of best fit to each replicate was made. The slope values for these linear lines indicated the speed of initial reaction. Each replicate was normalised to the original Con1 protease by division of the average slope value.

### Machine learning implementation

Best candidates for the second round of activity screening were generated by batch Bayesian optimisation using the Active Learning-Assisted Directed Evolution (ALDE) package [19].The training dataset consisted of the starting Con1 sequence and associated fitness score (1), plus 48 Con1 variants, randomly mutated at positions X, Y and Z with associated FRET activity used as a measure of fitness. Amino acids were represented using one-hot encoding. The surrogate model used to evaluate the fitness landscape over all possible amino acid combinations was an ensemble of 5 deep neural networks (DNN) with bootstrapping, where each DNN was supplied with random 90% of the available training data and treated as samples from a Bayesian posterior distribution. At each iteration of Bayesian optimisation, the proposal that maximised the Thomson Sampling acquisition function from a DNN randomly sampled from the ensemble which was then used as the input for the next iteration. After completion, the 32 top suggested variants were selected as candidates for a second round of FRET activity measurements. Computation was performed using the resources provided by the Edinburgh Compute and Data Facility (ECDF) (http://www.ecdf.ed.ac.uk/).

## Data availability

All data for this article is available in the supplementary information, alongside further supporting figures and text.

## Author information

### Affiliations

Centre for Engineering Biology, Institute of Quantitative Biology, Biochemistry and Biotechnology, School of Biological Sciences, University of Edinburgh, EH9 3BF, Scotland: Ella Lucille Thornton, Jeremy T. Boyle, Nadanai Laohakunakorn, Lynne Regan

### Contributions

ELT: Conceptualisation, investigation, analysis, writing - original draft, writing - editing JTB: Analysis, writing - editing

NL: Conceptualisation, supervision, writing - editing

LR: Conceptualisation, supervision, writing – original draft, writing - editing

## Acknowledgements

We thank Haresh Bhaskar, Hannah Johns, Kasia Stefaniak and Fokhrul Islam for valuable feedback on the manuscript. We thank Jason Yang and other members of the Arnold lab for technical support in running the ALDE code. We acknowledge the support of the School of Biological Sciences at the University of Edinburgh and the use of the resources provided by the Edinburgh Compute and Data Facility (ECDF) (http://www.ecdf.ed.ac.uk/). LR, ELT, & NL are grateful for the support of the Leverhulme Trust (RPG-2021-230). JTB & LR are grateful for the support of the Wellcome Trust via the Integrative Cell Mechanisms PhD program (218470/Z/19/Z). NL gratefully acknowledges the support of a UKRI Future Leaders Fellowship (MR/V027107/1).

