## Supplementary information for "Cell-free protein synthesis as a method to rapidly screen machine learning-directed protease variants"

Thornton *et al.*

Supplementary Figures:

1. DoE for protease expression
2. Calibration of FRET substrate against known concentrations of protease
3. Identity of all original random mutants
4. Time-course FRET data for 48 B mutants
5. Dataset of FRET data truncated at 0.55
6. Validation of fitness scoring against FRET time-course data
7. Full list of output variants from ALDE
8. Affinity purification SDS-PAGE
9. Size exclusion chromatography SDS-PAGE
10. Purified variants stocked at equimolar concentrations SDS-PAGE

Supplementary tables:

1. Con1 annotated DNA and protein sequence
2. FRET substrate annotated DNA and protein sequence
3. Oligonucleotides used in this study

Supplementary data (provided as individual files):

1. Figure 4 dataset A mutants
2. Figure 4 dataset B mutants
3. ML suggested variant dataset for Figure 5
4. Comparison of CFPS and purified variants for Figure 6

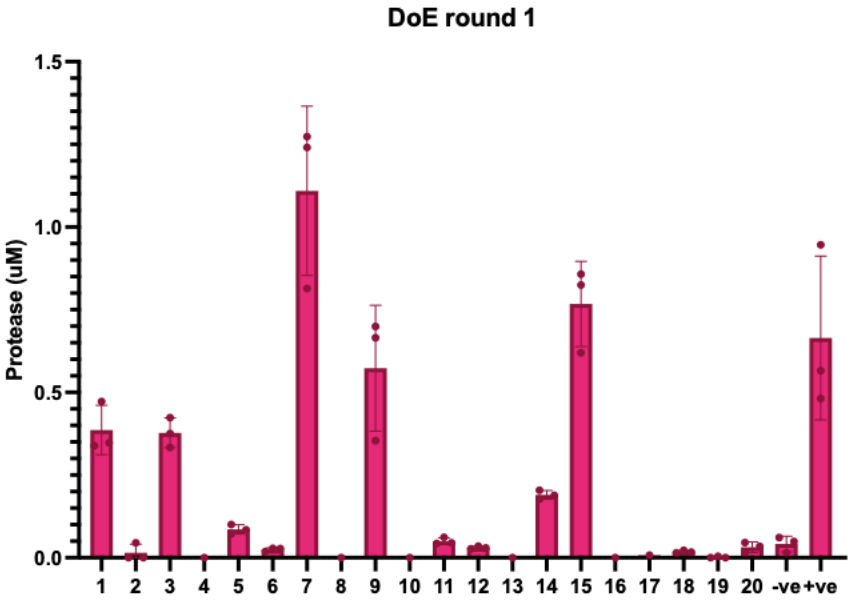


| First round: | | | |
|---|---|---|---|
| Total K (mM) | Total Mg (mM) | PEG (%) | Yield (uM) |
| 110.00 | 11.00 | 3 | 0.39 |
| 228.90 | 26.74 | 3 | 0.02 |
| 227.46 | 10.41 | 5 | 0.38 |
| 107.70 | 26.46 | 5 | 0.00 |
| 168.78 | 18.72 | 4 | 0.09 |
| 168.78 | 18.72 | 4 | 0.03 |
| 229.86 | 10.97 | 3 | 1.11 |
| 110.00 | 27.00 | 3 | 0.00 |
| 108.66 | 10.69 | 5 | 0.57 |
| 226.50 | 26.18 | 5 | 0.00 |
| 168.78 | 18.72 | 4 | 0.05 |
| 168.78 | 18.72 | 4 | 0.03 |
| 94.53 | 18.89 | 4 | 0.00 |
| 243.03 | 18.54 | 4 | 0.19 |
| 169.38 | 8.86 | 4 | 0.77 |
| 168.18 | 28.58 | 4 | 0.00 |
| 170.00 | 19.00 | 2.75 | 0.01 |
| 167.28 | 18.37 | 5.25 | 0.02 |
| 168.78 | 18.72 | 4 | 0.00 |
| 168.78 | 18.72 | 4 | 0.03 |

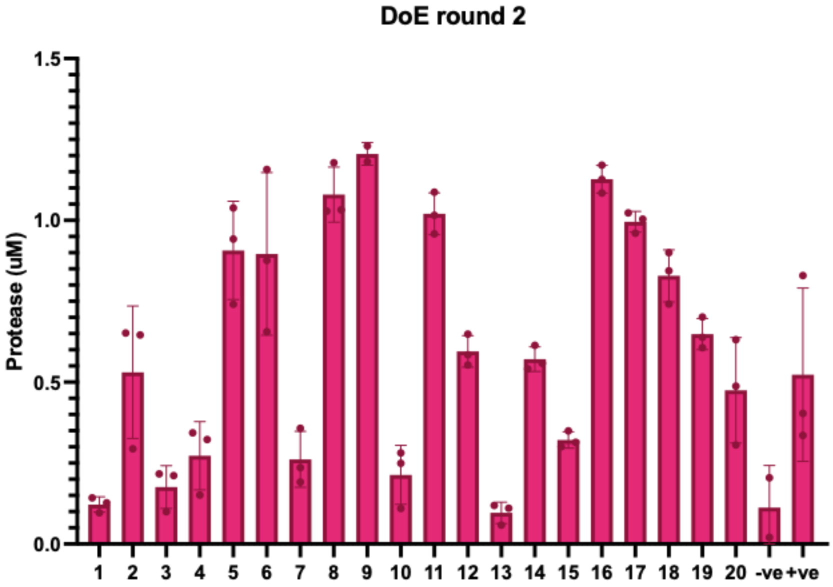


| Second round: | | | |
|---|---|---|---|
| K | Mg | PEG | Yield |
| 220 | 7.7 | 1 | 0.12 |
| 300 | 12 | 1 | 0.53 |
| 299.36 | 7.5502 | 3 | 0.18 |
| 219.901 | 11.97 | 3 | 0.27 |
| 270 | 10.5 | 2 | 0.91 |
| 270 | 10.5 | 2 | 0.90 |
| 300 | 7.7 | 1 | 0.26 |
| 220 | 12 | 1 | 1.08 |
| 220 | 7.7 | 3 | 1.21 |
| 299.1 | 11.79 | 3 | 0.21 |
| 270 | 10.5 | 2 | 1.02 |
| 270 | 10.5 | 2 | 0.60 |
| 207.5 | 10.5 | 2 | 0.10 |
| 307.5 | 10.5 | 2 | 0.57 |
| 270 | 7 | 2 | 0.32 |
| 270 | 12.375 | 2 | 1.13 |
| 270 | 10.5 | 0.75 | 1.00 |
| 269.1 | 10.29 | 3.25 | 0.83 |
| 270 | 10.5 | 2 | 0.65 |
| 270 | 10.5 | 2 | 0.47 |

**Figure S1. A Design of Experiments approach to optimise protease expression in CFPS**. The experimental design of Central Composite Design was utilised, with axial points. Mg, K, and PEG were selected as variables to assess. The range of concentrations to be tested for each was decided by a literature review. The amount of protease made in each condition was assessed by an activity assay and compared to a purified calibration of protease against FRET substrate (S2). Round one identified conditions that were more favourable for protease expression (as seen by comparison with the +ve sample, with original energy solution components). To further optimise the conditions, another round of DoE was planned and completed, finding a condiiton that increased protease expression from ~0.5 μM to ~1 μM. These conditions were used for all subsequent CFPS experiments.

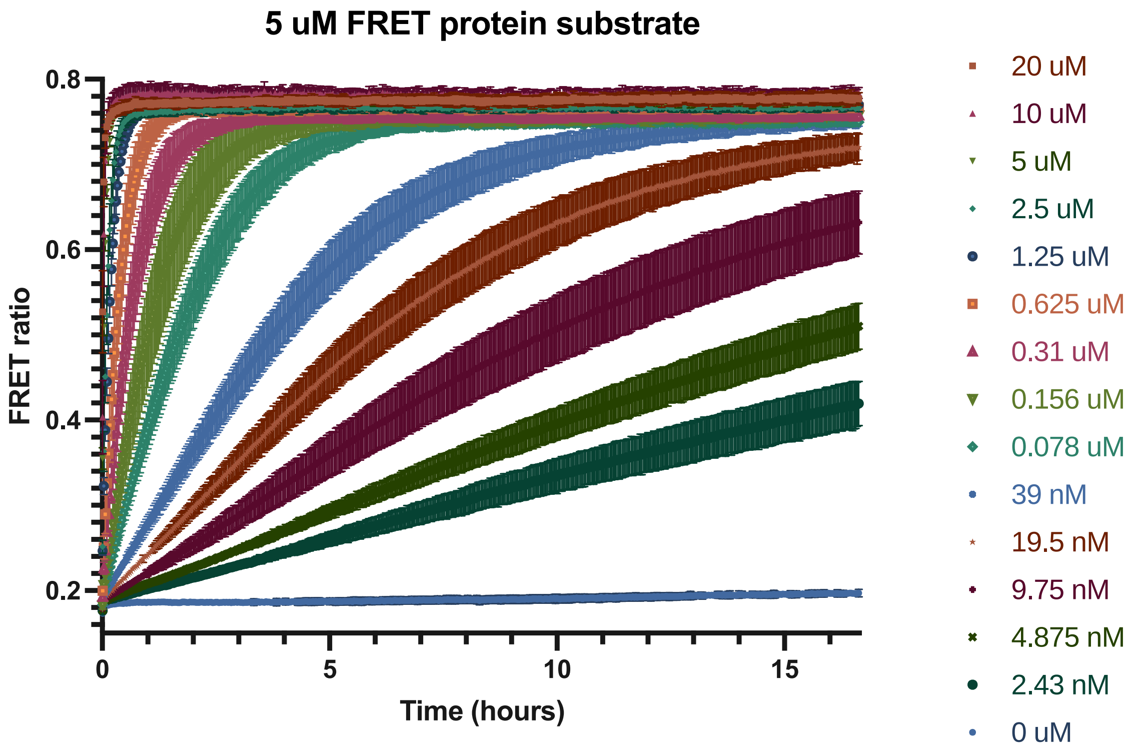


**Figure S2. Calibration of FRET substrate against known concentrations of purified Con1 protease.** FRET substrate was mixed together with protease in buffer, to give a final FRET concentration of 5 μM. This was immediately measured in a plate reader for fluorescence and subsequent assessment of FRET over time.

### Region A:

| | Residue: | | |
|---|---|---|---|
| Mutant no: | 167 | 169 | 172 |
| 1 | E | I | N |
| 2 | T | H | T |
| 3 | R | M | F |
| 4 | L | L | A |
| 5 | R | R | L |
| 6 | W | A | K |
| 7 | R | K | L |
| 8 | W | A | F |
| 9 | G | F | F |
| 10 | Q | L | F |
| 11 | H | T | V |
| 12 | G | Y | R |
| 13 | V | M | N |
| 14 | V | T | D |
| 15 | Y | F | N |
| 16 | W | L | G |
| 17 | Q | R | E |
| 18 | E | R | W |
| 19 | M | S | I |
| 20 | D | L | G |
| 21 | E | D | W |
| 22 | K | S | F |
| 23 | K | I | W |
| 24 | D | A | G |
| 25 | G | S | Y |
| 26 | L | T | K |
| 27 | R | A | H |
| 28 | H | M | E |
| 29 | Q | V | T |
| 30 | N | K | D |
| 31 | E | H | S |
| 32 | S | R | F |
| 33 | W | R | E |
| 34 | A | L | F |
| 35 | S | W | S |
| 36 | I | E | N |
| 37 | I | L | H |
| 38 | L | M | D |
| 39 | I | A | N |
| 40 | L | M | F |
| 41 | I | Y | I |
| 42 | S | T | E |
| 43 | D | D | T |
| 44 | E | Q | K |
| 45 | N | Q | Y |
| 46 | S | D | R |
| 47 | Q | L | G |
| 48 | K | D | I |

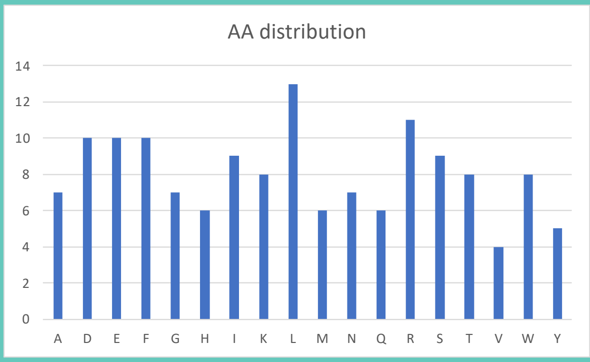


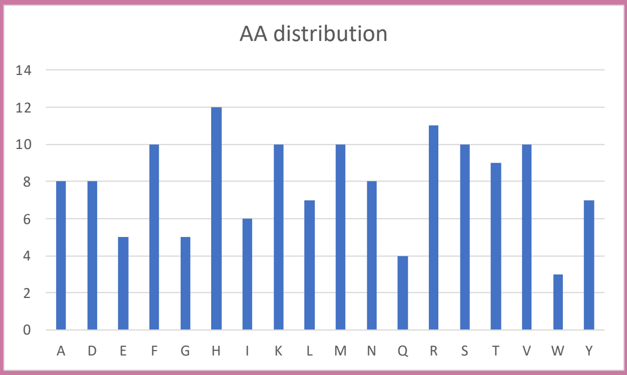


### Region B:

| | Residue: | | |
|---|---|---|---|
| Mutant no: | 217 | 218 | 219 |
| 1 | H | V | L |
| 2 | Y | L | I |
| 3 | E | G | K |
| 4 | N | N | R |
| 5 | Q | T | H |
| 6 | R | M | H |
| 7 | W | A | K |
| 8 | H | S | Y |
| 9 | H | M | Y |
| 10 | L | N | M |
| 11 | A | M | G |
| 12 | K | F | W |
| 13 | F | H | S |
| 14 | S | H | F |
| 15 | A | V | E |
| 16 | M | D | H |
| 17 | I | V | T |
| 18 | S | K | Q |
| 19 | D | H | R |
| 20 | R | S | K |
| 21 | I | E | D |
| 22 | A | R | V |
| 23 | W | R | H |
| 24 | A | R | F |
| 25 | M | D | Y |
| 26 | S | V | S |
| 27 | G | Y | Y |
| 28 | G | N | H |
| 29 | F | R | F |
| 30 | R | T | A |
| 31 | T | K | K |
| 32 | F | S | L |
| 33 | R | K | N |
| 34 | M | T | F |
| 35 | Y | G | D |
| 36 | K | M | I |
| 37 | A | K | T |
| 38 | F | L | T |
| 39 | Y | V | D |
| 40 | F | L | V |
| 41 | I | E | N |
| 42 | V | A | N |
| 43 | M | Q | H |
| 44 | I | S | D |
| 45 | E | V | T |
| 46 | N | Q | V |
| 47 | S | D | T |
| 48 | M | R | L |

**Figure S3. Identity of all random mutants.** 48 random mutants were created for both region A and region B by a script which compiled a randomised listed of amino acids into a 3 x 48 table. At this stage of the design process, we removed proline and cysteine as possibilities for mutagenesis. The representation of all other amino acids was verified as seen in the bar plots.

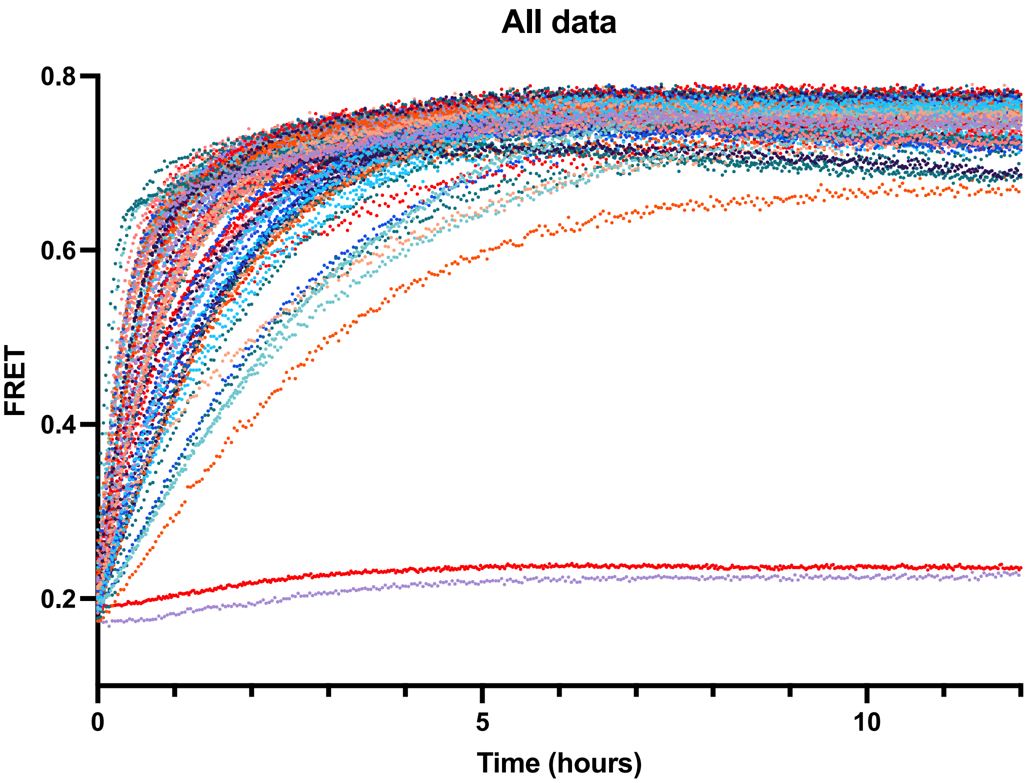


**Figure S4. Example time-course data of FRET for 48 B mutants after production by CFPS.**

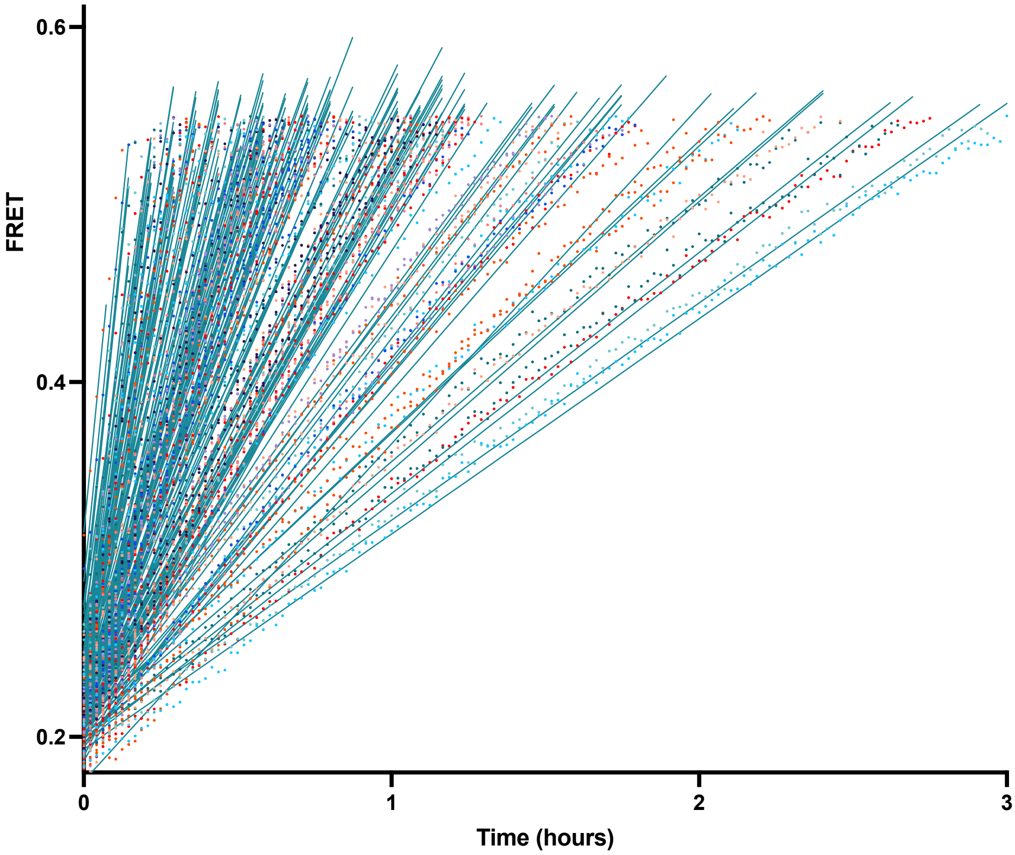


**Figure S5. Data from S4 truncated at FRET = 0.55.** Plot is cropped to Time = 3 hours to allow closer examination of data points and linear fit. We found this data processing step allowed for accurate scoring of mutant activity, as shown further in Figure S6.

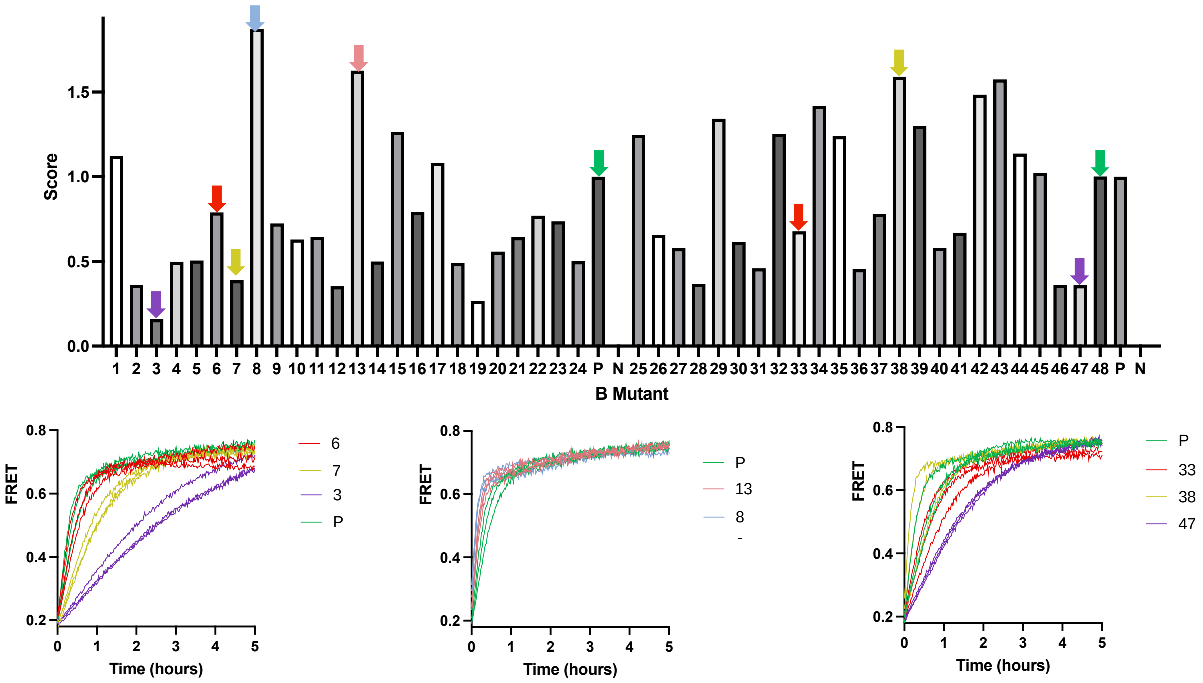


**Figure S6. Assessment of final fitness score accuracy against original FRET time-course data.** Final scores (average of 3 replicates) are shown in the bar chart for all region B mutants. Below, line charts show individual replicates for selected mutants, the colour of which is then matched to their identity and to the corresponding bar.

**Region A:**

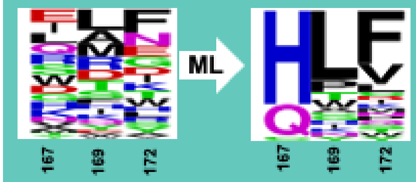


| Mutant: | 167 | 169 | 172 |
|---|---|---|---|
| 1 | H | L | Q |
| 2 | H | L | P |
| 3 | H | L | C |
| 4 | H | P | F |
| 5 | H | L | D |
| 6 | Q | L | V |
| 7 | H | L | W |
| 8 | H | L | L |
| 9 | H | L | Y |
| 10 | H | L | K |
| 11 | H | K | F |
| 12 | H | L | N |
| 13 | H | L | S |
| 14 | Q | T | V |
| 15 | Q | L | L |
| 16 | H | Y | F |
| 17 | H | L | M |
| 18 | Q | T | F |
| 19 | H | I | F |
| 20 | Q | P | F |
| 21 | H | L | I |
| 22 | H | L | T |
| 23 | Q | W | V |
| 24 | C | L | F |
| 25 | H | L | V |
| 26 | H | C | F |
| 27 | H | H | F |
| 28 | H | N | F |
| 29 | H | G | F |
| 30 | H | A | F |
| 31 | Q | P | V |
| 32 | Q | W | F |
| Con1 | H | L | F |

**Region B:**

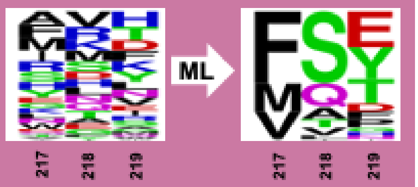


| Mutant: | 217 | 218 | 219 |
|---|---|---|---|
| 1 | V | S | D |
| 2 | F | S | Y |
| 3 | F | S | T |
| 4 | F | T | T |
| 5 | V | S | Y |
| 6 | F | Q | E |
| 7 | F | T | Y |
| 8 | M | S | Y |
| 9 | F | A | Y |
| 10 | V | S | T |
| 11 | F | S | P |
| 12 | V | Q | T |
| 13 | M | Q | Y |
| 14 | F | S | E |
| 15 | F | V | E |
| 16 | F | A | T |
| 17 | V | S | E |
| 18 | M | Q | E |
| 19 | M | S | T |
| 20 | F | A | E |
| 21 | F | Q | Y |
| 22 | V | S | F |
| 23 | M | S | E |
| 24 | F | S | D |
| 25 | V | Q | Y |
| 26 | F | S | H |
| 27 | F | S | N |
| 28 | F | S | G |
| 29 | F | S | F |
| 30 | M | S | P |
| 31 | F | L | E |
| 32 | M | V | E |
| Con1 | L | Q | E |

**Figure S7. Identity of mutants suggested by ALDE workflow.** Sequence logos of the amino acid identities of the random mutants, compared with machine learning suggested mutants are shown. For region A, only the first 10 of these suggested mutants were tested. For region B, all 32 were screened. The identity of each variant at the residues to be mutated is indicated in the table, with amino acids coloured based on type: green = hydrophobic, pink = polar, red = negatively charged, orange = small non-polar, blue = positively charged.

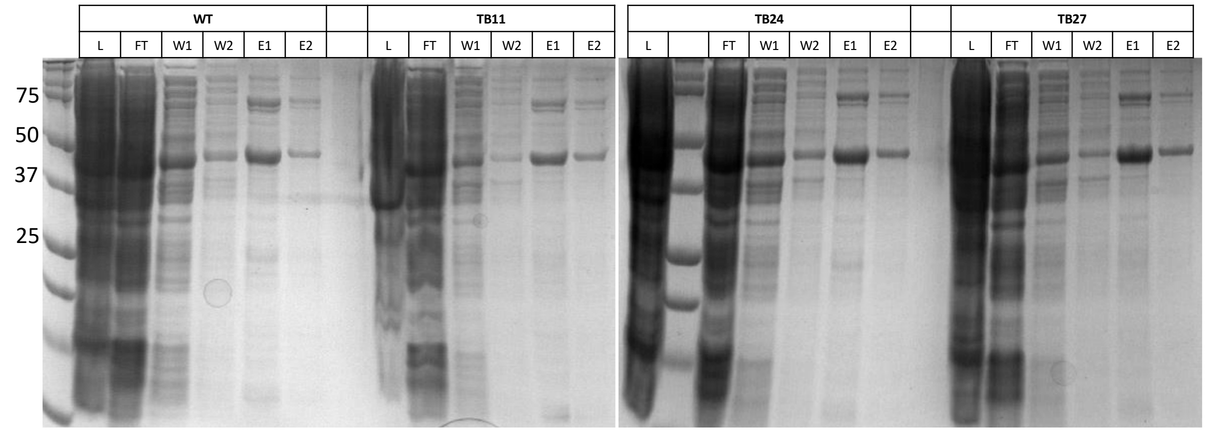


**Figure S8. Affinity purification of four protease variants.** Indicated by the lane title, different samples from the purification process were loaded: L (lysate), FT (flow-through), W1 (wash 1), W2 (wash 2), E1 (elution 1), E2 (elution 2). Expected MW = 42 kDa.

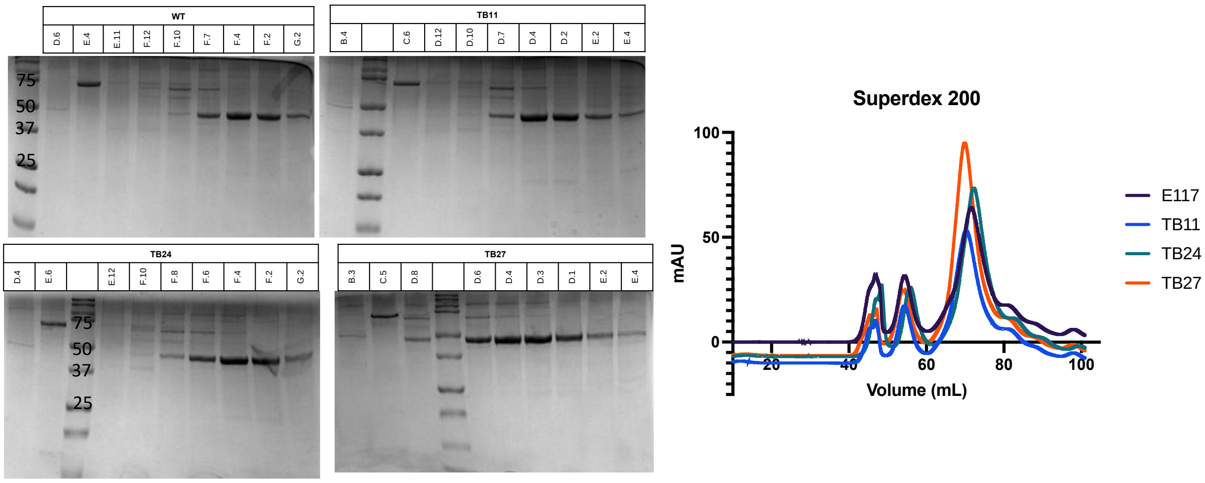


**Figure S9. Size exclusion chromatography of four protease variants.** Fractions eluted from the column were run on SDS-PAGE to assess purity. Absorbance from the SEC run is shown on the line plot. The largest peak contained the proteases.

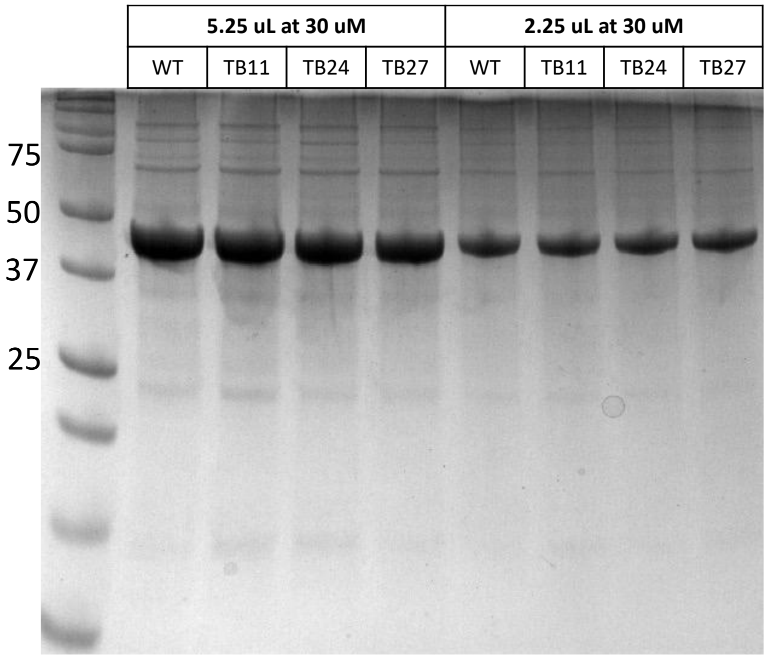


**Figure S10. Purified variants stocked at equimolar concentrations.**

| **Original Con1** | |
|---|---|
| DNA sequence | ATGCACCATCACCATCACCATGCCGTAACCACCTTATCAGGTTTATCAGGTGA<br>GCAAGGTCCGTCCGGTGATATGACAACTGAAGAAGATAGTGCTACCCATATT<br>AAATTCTCAAAACGTGATGAGGACGGCCGTGAGTTAGCTGGTGCAACTATG<br>GAGTTGCGTGATTCATCTGGTAAAACTATTAGTACATGGATTTCAGATGGACA<br>TGTGAAGGATTTCTACCTGTATCCAGGAAAATATACATTTGTCGAAACCGCAG<br>CACCAGACGGTTATGAGGTAGCAACTCCAATTGAATTTACAGTTAATGAGGA<br>CGGTCAGGTTACTGTAGATGGTGAAGCAACTGAAGGTGACGCTCATACTGG<br>AGGTAGCGGTGGCTCAAGCAAATCTTTATTCCGTGGATTACGCGATTACAAC<br>CCGATCGCCAGCAATATTTGTCATTTGACTAATGAGAGCGACGGTCATAGTAA<br>CTCGTTGTATGGCATCGGGTTTGGGCCGCTTATTATTACAAACCAACATTTGT<br>TTCGTCGCAACAACGGGGAACTTACGATTCAATCTCGCCATGGCGAGTTCG<br>TAGTTAAGAACACCACGCAACTTAAGCTTCTGCCAATCGATGGGCGTGATAT<br>CCTTATCATTCGTTTGCCCAAAGATTTCCCACCCTTTCCACAGAAACTTAAAT<br>TCCGCCAACCTGAAAAAGGCGAACGCATTTGCTTGGTTGGCTCGAATTTTCA<br>GACCAAGTCCATCACCAGTACAGTCAGCGAAACATCAACAACCATGCCGGT<br>GGAGAATAGTCAATTTTGGAAGCACTGGATTTCAACCAAAGACGGTCATTGT<br>GGCCTGCCTCTTGTTTCCACGAAGGATGGCAAGATCCTGGGCATCCATTCTT<br>TGGCGAATTTTACCAATACGATCAACTACTTCGCCGCATTCCCAGAAGATTTT<br>GAGGAGACGTATCTTCATACCCAAGAAGCTCAAGAATGGGTAAAGCACTGGA<br>AATATAACCCGGATGCGATCTCCTGGGGGAGCCTTAACCTTCAAGAAAGCCA<br>ACCAGAAGAGCCTTTCAAAATCGTAAAATTAGTAACGGACGGAGGTAGCGGT<br>GGCTCAGAACAGAAACTGATTAGCGAAGAAGATCTGTAA |
| Protein sequence | MHHHHHHAVTTLSGLSGEQGPSGDMTTEEDSATHIKFSKRDEDGRELAGATM<br>ELRDSSGKTISTWISDGHVKDFYLYPGKYTFVETAAPDGYEVATPIEFTVNEDG<br>QVTVDGEATEGDAHTGGSGGSSKSLFRGLRDYNPIASNICHLTNESDGHSNSL<br>YGIGFGPLIITNQHLFRRNNGELTIQSRHGEFVVKNTTQLKLLPIDGRDILIIRLPK<br>DFPPFPQKLKFRQPEKGERICLVGSNFQTKSITSTVSETSTTMPVENSQFWKH<br>WISTKDGHCGLPLVSTKDGKILGIHSLANFTNTINYFAAFPEDFEETYLHTQEAQ<br>EWVKHWKYNPDAISWGSLNLQESQPEEPFKIVKLVTDGGSGGSEQKLISEEDL<br>* |
| Identity | His-SpyCatcher-Con1-myc |

Table S1. Annotated Con1 DNA and protein sequence.

| **FRET substrate** | |
|---|---|
| DNA sequence | ATGCGGGGTTCTCATCATCATCATCATCATGGTATGGCTAGCATGACTGGTG<br>GACAGCAAATGGGTCGGGATCTGTACGACGATGACGATAAGGATCCGGGCC<br>GCATGGTGAGCAAGGGCGAGGAGCTGTTCACCGGGGTGGTGCCCATCCTG<br>GTCGAGCTGGACGGCGACGTAAACGGCCACAAGTTCAGCGTGTCCGGCGA<br>GGGCGAGGGCGATGCCACCTACGGCAAGCTGACCCTGAAGTTCATCTGCA<br>CCACCGGCAAGCTGCCCGTGCCCTGGCCCACCCTCGTGACCACCCTGACC<br>TGGGGCGTGCAGTGCTTCAGCCGCTACCCCGACCACATGAAGCAGCACGA<br>CTTCTTCAAGTCCGCCATGCCCGAAGGCTACGTCCAGGAGCGCACCATCTT<br>CTTCAAGGACGACGGCAACTACAAGACCCGCGCCGAGGTGAAGTTCGAGG<br>GCGACACCCTGGTGAACCGCATCGAGCTGAAGGGCATCGACTTCAAGGAG<br>GACGGCAACATCCTGGGGCACAAGCTGGAGTACAACTACATCAGCCACAAC<br>GTCTATATCACCGCCGACAAGCAGAAGAACGGCATCAAGGCCAACTTCAAG<br>ATCCGCCACAACATCGAGGACGGCAGCGTGCAGCTCGCCGACCACTACCA<br>GCAGAACACCCCCATCGGCGACGGCCCCGTGCTGCTGCCCGACAACCACT<br>ACCTGAGCACCCAGTCCGCCCTGAGCAAAGACCCCAACGAGAAGCGCGAT<br>CACATGGTCCTGCTGGAGTTCGTGACCGCCGCCGGGATCGGTACCGTAGG<br>ATTTCTAACAGCGACCGAAGCAGTGTATCATCAATCCCTGGATTCCATCACC<br>GTTAACGGTACCGGTGGAATGGTGAGCAAGGGCGAGGAGCTGTTCACCGG |

| | |
|---|---|
| | GGTGGTGCCCATCCTGGTCGAGCTGGACGGCGACGTAAACGGCCACAAGT<br>TCAGCGTGTCCGGCGAGGGCGAGGGCGATGCCACCTACGGCAAGCTGAC<br>CCTGAAGTTCATCTGCACCACCGGCAAGCTGCCCGTGCCCTGGCCCACCC<br>TCGTGACCACCTTCGGCTACGGCCTGCAGTGCTTCGCCCGCTACCCCGAC<br>CACATGAAGCAGCACGACTTCTTCAAGTCCGCCATGCCCGAAGGCTACGTC<br>CAGGAGCGCACCATCTTCTTCAAGGACGACGGCAACTACAAGACCCGCGC<br>CGAGGTGAAGTTCGAGGGCGACACCCTGGTGAACCGCATCGAGCTGAAGG<br>GCATCGACTTCAAGGAGGACGGCAACATCCTGGGGCACAAGCTGGAGTAC<br>AACTACAACAGCCACAACGTCTATATCATGGCCGACAAGCAGAAGAACGGCA<br>TCAAGGTGAACTTCAAGATCCGCCACAACATCGAGGACGGCAGCGTGCAG<br>CTCGCCGACCACTACCAGCAGAACACCCCCATCGGCGACGGCCCCGTGCT<br>GCTGCCCGACAACCACTACCTGAGCTACCAGTCCGCCCTGAGCAAAGACC<br>CCAACGAGAAGCGCGATCACATGGTCCTGCTGGAGTTCGTGACCGCCGCC<br>GGGATCACTCTCGGCATGGACGAGCTGTACAAGTAA |
| Protein sequence | MRGSHHHHHHGMASMTGGQQMGRDLYDDDDKDPGRMVSKGEELFTGVVPIL<br>VELDGDVNGHKFSVSGEGEGDATYGKLTLKFICTTGKLPVPWPTLVTTLTWGV<br>QCFSRYPDHMKQHDFFKSAMPEGYVQERTIFFKDDGNYKTRAEVKFEGDTLV<br>NRIELKGIDFKEDGNILGHKLEYNYISHNVYITADKQKNGIKANFKIRHNIEDGSV<br>QLADHYQQNTPIGDGPVLLPDNHYLSTQSALSKDPNEKRDHMVLLEFVTAAGI<br>GTVGFLTATEAVYHQSLDSITVNGTGGMVSKGEELFTGVVPILVELDGDVNGHK<br>FSVSGEGEGDATYGKLTLKFICTTGKLPVPWPTLVTTFGYGLQCFARYPDHMK<br>QHDFFKSAMPEGYVQERTIFFKDDGNYKTRAEVKFEGDTLVNRIELKGIDFKED<br>GNILGHKLEYNYNSHNVYIMADKQKNGIKVNFKIRHNIEDGSVQLADHYQQNTPI<br>GDGPVLLPDNHYLSYQSALSKDPNEKRDHMVLLEFVTAAGITLGMDELYK* |
| Identity | His-CFP-cut site-YFP |

Table S2. Annotated FRET substrate DNA and protein sequence.

| Code | Purpose | Sequence |
|---|---|---|
| P291 | B_1_F | TTAACCATGTGCTGAGCCAACCAGAAGAGCCTT |
| P292 | B_1_R | GGCTCAGCACATGGTTAAGGCTCCCCCAGGA |
| P293 | A_1_F | GAATCGATTGCCAACAACACCAATACGATCAACTACT |
| P294 | A_1_R | GTTGTTGGCAATCGATTCGATGCCCAGGATCT |
| P298 | B_2_F | TTAACTATCTGATTAGCCAACCAGAAGAGCCTT |
| P299 | B_2_R | GGCTAATCAGATAGTTAAGGCTCCCCCAGGA |
| P300 | B_3_F | TTAACGAAGGCAAAAGCCAACCAGAAGAGCCTT |
| P301 | B_3_R | GGCTTTTGCCTTCGTTAAGGCTCCCCCAGGA |
| P302 | B_4_F | TTAACAACAATCGCAGCCAACCAGAAGAGCCTT |
| P303 | B_4_R | GGCTGCGATTGTTGTTAAGGCTCCCCCAGGA |
| P304 | B_5_F | TTAACCAGACCCATAGCCAACCAGAAGAGCCTT |
| P305 | B_5_R | GGCTATGGGTCTGGTTAAGGCTCCCCCAGGA |
| P306 | B_6_F | TTAACCGCATGCATAGCCAACCAGAAGAGCCTT |
| P307 | B_6_R | GGCTATGCATGCGGTTAAGGCTCCCCCAGGA |
| P308 | B_7_F | TTAACTGGGCGAAAAGCCAACCAGAAGAGCCTT |
| P309 | B_7_R | GGCTTTTCGCCCAGTTAAGGCTCCCCCAGGA |
| P310 | B_8_F | TTAACCATAGCTATAGCCAACCAGAAGAGCCTT |
| P311 | B_8_R | GGCTATAGCTATGGTTAAGGCTCCCCCAGGA |
| P312 | B_9_F | TTAACCATATGTATAGCCAACCAGAAGAGCCTT |
| P313 | B_9_R | GGCTATACATATGGTTAAGGCTCCCCCAGGA |
| P314 | B_10_F | TTAACCTGAACATGAGCCAACCAGAAGAGCCTT |
| P315 | B_10_R | GGCTCATGTTCAGGTTAAGGCTCCCCCAGGA |
| P317 | B_11_R | GGCTGCCCATCGCGTTAAGGCTCCCCCAGGA |
| P318 | B_12_F | TTAACAAATTTTGGAGCCAACCAGAAGAGCCTT |
| P319 | B_12_R | GGCTCCAAAATTTGTTAAGGCTCCCCCAGGA |
| P320 | B_13_F | TTAACTTTCATAGCAGCCAACCAGAAGAGCCTT |
| P321 | B_13_R | GGCTGCTATGAAAGTTAAGGCTCCCCCAGGA |
| P322 | B_14_F | TTAACAGCCATTTTAGCCAACCAGAAGAGCCTT |
| P323 | B_14_R | GGCTAAAATGGCTGTTAAGGCTCCCCCAGGA |
| P325 | B_15_R | GGCTTTCCACCGCGTTAAGGCTCCCCCAGGA |
| P326 | B_16_F | TTAACATGGATCATAGCCAACCAGAAGAGCCTT |
| P327 | B_16_R | GGCTATGATCCATGTTAAGGCTCCCCCAGGA |
| P328 | B_17_F | TTAACATTGTGACCAGCCAACCAGAAGAGCCTT |
| P329 | B_17_R | GGCTGGTCACAATGTTAAGGCTCCCCCAGGA |
| P330 | B_18_F | TTAACAGCAAACAGAGCCAACCAGAAGAGCCTT |

| | | |
|---|---|---|
| P331 | B_18_R | GGCTCTGTTTGCTGTTAAGGCTCCCCCAGGA |
| P332 | B_19_F | TTAACGATCATCGCAGCCAACCAGAAGAGCCTT |
| P333 | B_19_R | GGCTGCGATGATCGTTAAGGCTCCCCCAGGA |
| P334 | B_20_F | TTAACCGCAGCAAAAGCCAACCAGAAGAGCCTT |
| P335 | B_20_R | GGCTTTTGCTGCGGTTAAGGCTCCCCCAGGA |
| P336 | B_21_F | TTAACATTGAAGATAGCCAACCAGAAGAGCCTT |
| P337 | B_21_R | GGCTATCTTCAATGTTAAGGCTCCCCCAGGA |
| P341 | B_23_R | GGCTATGGCGCCAGTTAAGGCTCCCCCAGGA |
| P343 | B_24_R | GGCTAAAGCGCGCGTTAAGGCTCCCCCAGGA |
| P344 | B_25_F | TTAACATGGATTATAGCCAACCAGAAGAGCCTT |
| P345 | B_25_R | GGCTATAATCCATGTTAAGGCTCCCCCAGGA |
| P347 | B_26_R | GGCTGCTCACGCTGTTAAGGCTCCCCCAGGA |
| P348 | B_27_F | TTAACGGCTATTATAGCCAACCAGAAGAGCCTT |
| P349 | B_27_R | GGCTATAATAGCCGTTAAGGCTCCCCCAGGA |
| P350 | B_28_F | TTAACGGCAACCATAGCCAACCAGAAGAGCCTT |
| P351 | B_28_R | GGCTATGGTTGCCGTTAAGGCTCCCCCAGGA |
| P352 | B_29_F | TTAACTTTCGCTTTAGCCAACCAGAAGAGCCTT |
| P353 | B_29_R | GGCTAAAGCGAAAGTTAAGGCTCCCCCAGGA |
| P354 | B_30_F | TTAACCGCACCGCGAGCCAACCAGAAGAGCCTT |
| P355 | B_30_R | GGCTCGCGGTGCGGTTAAGGCTCCCCCAGGA |
| P356 | B_31_F | TTAACACCAAAAAGAGCCAACCAGAAGAGCCTT |
| P357 | B_31_R | GGCTCTTTTTGGTGTTAAGGCTCCCCCAGGA |
| P358 | B_32_F | TTAACTTTAGCCTGAGCCAACCAGAAGAGCCTT |
| P359 | B_32_R | GGCTCAGGCTAAAGTTAAGGCTCCCCCAGGA |
| P360 | B_33_F | TTAACCGCAAAAACAGCCAACCAGAAGAGCCTT |
| P361 | B_33_R | GGCTGTTTTTGCGGTTAAGGCTCCCCCAGGA |
| P362 | B_34_F | TTAACATGACCTTCAGCCAACCAGAAGAGCCTT |
| P363 | B_34_R | GGCTGAAGGTCATGTTAAGGCTCCCCCAGGA |
| P364 | B_35_F | TTAACTATGGCGATAGCCAACCAGAAGAGCCTT |
| P365 | B_35_R | GGCTATCGCCATAGTTAAGGCTCCCCCAGGA |
| P366 | B_36_F | TTAACAAAATGATTAGCCAACCAGAAGAGCCTT |
| P367 | B_36_R | GGCTAATCATTTTGTTAAGGCTCCCCCAGGA |
| P368 | B_37_F | TTAACGCGAAAACCAGCCAACCAGAAGAGCCTT |
| P369 | B_37_R | GGCTGGTTTTCGCGTTAAGGCTCCCCCAGGA |
| P370 | B_38_F | TTAACTTCCTGACCAGCCAACCAGAAGAGCCTT |
| P371 | B_38_R | GGCTGGTCAGGAAGTTAAGGCTCCCCCAGGA |
| P372 | B_39_F | TTAACTATGTGGATAGCCAACCAGAAGAGCCTT |
| P373 | B_39_R | GGCTATCCACATAGTTAAGGCTCCCCCAGGA |
| P374 | B_40_F | TTAACTTCCTGGTGAGCCAACCAGAAGAGCCTT |
| P375 | B_40_R | GGCTCACCAGGAAGTTAAGGCTCCCCCAGGA |
| P376 | B_41_F | TTAACATTGAAAACAGCCAACCAGAAGAGCCTT |
| P377 | B_41_R | GGCTGTTTTCAATGTTAAGGCTCCCCCAGGA |
| P378 | B_42_F | TTAACGTGGCGAACAGCCAACCAGAAGAGCCTT |
| P379 | B_42_R | GGCTGTTCGCCACGTTAAGGCTCCCCCAGGA |
| P380 | B_43_F | TTAACATGCAGCATAGCCAACCAGAAGAGCCTT |
| P381 | B_43_R | GGCTATGCTGCATGTTAAGGCTCCCCCAGGA |
| P382 | B_44_F | TTAACATTAGCGATAGCCAACCAGAAGAGCCTT |
| P383 | B_44_R | GGCTATCGCTAATGTTAAGGCTCCCCCAGGA |
| P384 | B_45_F | TTAACGAAGTGACCAGCCAACCAGAAGAGCCTT |
| P385 | B_45_R | GGCTGGTCACTTCGTTAAGGCTCCCCCAGGA |
| P386 | B_46_F | TTAACAACCAGGTGAGCCAACCAGAAGAGCCTT |
| P387 | B_46_R | GGCTCACCTGGTTGTTAAGGCTCCCCCAGGA |
| P388 | B_47_F | TTAACAGCGATACCAGCCAACCAGAAGAGCCTT |
| P389 | B_47_R | GGCTGGTATCGCTGTTAAGGCTCCCCCAGGA |
| P390 | B_48_F | TTAACATGCGCCTGAGCCAACCAGAAGAGCCTT |
| P391 | B_48_R | GGCTCAGGCGCATGTTAAGGCTCCCCCAGGA |
| P392 | A_2_F | ACCTCGCATGCCAACACCACCAATACGATCAACTACT |
| P393 | A_2_R | GGTGTTGGCATGCGAGGTGATGCCCAGGATCT |
| P394 | A_3_F | CGTTCGATGGCCAACTTTACCAATACGATCAACTACT |
| P395 | A_3_R | AAAGTTGGCCATCGAACGGATGCCCAGGATCT |
| P396 | A_4_F | CTGTCGCTGGCCAACGCGACCAATACGATCAACTACT |
| P397 | A_4_R | CGCGTTGGCCAGCGACAGGATGCCCAGGATCT |
| P398 | A_5_F | CGCTCGCGCGCCAACCTGACCAATACGATCAACTACT |
| P399 | A_5_R | CAGGTTGGCGCGCGAGCGGATGCCCAGGATCT |
| P400 | A_6_F | TGGTCGGCGGCCAACAAAACCAATACGATCAACTACT |
| P401 | A_6_R | TTTGTTGGCCGCCGACCAGATGCCCAGGATCT |
| P402 | A_7_F | CGCTCGAAAGCCAACCTGACCAATACGATCAACTACT |
| P403 | A_7_R | CAGGTTGGCTTTCGAGCGGATGCCCAGGATCT |
| P404 | A_8_F | TGGTCGGCGGCCAACTTTACCAATACGATCAACTACT |
| P405 | A_8_R | AAAGTTGGCCGCCGACCAGATGCCCAGGATCT |

| | | |
|---|---|---|
| P406 | A_9_F | GGCTCGTTTGCCAACTTTACCAATACGATCAACTACT |
| P407 | A_9_R | AAAGTTGGCAAACGAGCCGATGCCCAGGATCT |
| P408 | A_10_F | CAGTCGCTGGCCAACTTTACCAATACGATCAACTACT |
| P409 | A_10_R | AAAGTTGGCCAGCGACTGGATGCCCAGGATCT |
| P410 | A_11_F | CATTCGACCGCCAACGTGACCAATACGATCAACTACT |
| P411 | A_11_R | CACGTTGGCGGTCGAATGGATGCCCAGGATCT |
| P412 | A_12_F | GGCTCGTATGCCAACCGCACCAATACGATCAACTACT |
| P413 | A_12_R | GCGGTTGGCATACGAGCCGATGCCCAGGATCT |
| P414 | A_13_F | GTGTCGATGGCCAACAACACCAATACGATCAACTACT |
| P415 | A_13_R | GTTGTTGGCCATCGACACGATGCCCAGGATCT |
| P416 | A_14_F | GTGTCGACCGCCAACGATACCAATACGATCAACTACT |
| P417 | A_14_R | ATCGTTGGCGGTCGACACGATGCCCAGGATCT |
| P418 | A_15_F | TATTCGTTTGCCAACAACACCAATACGATCAACTACT |
| P419 | A_15_R | GTTGTTGGCAAACGAATAGATGCCCAGGATCT |
| P420 | A_16_F | TGGTCGCTGGCCAACGGCACCAATACGATCAACTACT |
| P421 | A_16_R | GCCGTTGGCCAGCGACCAGATGCCCAGGATCT |
| P422 | A_17_F | CAGTCGCGCGCCAACGAAACCAATACGATCAACTACT |
| P423 | A_17_R | TTCGTTGGCGCGCGACTGGATGCCCAGGATCT |
| P424 | A_18_F | GAATCGCGCGCCAACTGGACCAATACGATCAACTACT |
| P425 | A_18_R | CCAGTTGGCGCGCGATTCGATGCCCAGGATCT |
| P426 | A_19_F | ATGTCGAGCGCCAACATTACCAATACGATCAACTACT |
| P427 | A_19_R | AATGTTGGCGCTCGACATGATGCCCAGGATCT |
| P428 | A_20_F | GATTCGCTGGCCAACGGCACCAATACGATCAACTACT |
| P429 | A_20_R | GCCGTTGGCCAGCGAATCGATGCCCAGGATCT |
| P430 | A_21_F | GAATCGGATGCCAACTGGACCAATACGATCAACTACT |
| P431 | A_21_R | CCAGTTGGCATCCGATTCGATGCCCAGGATCT |
| P432 | A_22_F | AAATCGAGCGCCAACTTTACCAATACGATCAACTACT |
| P433 | A_22_R | AAAGTTGGCGCTCGATTTGATGCCCAGGATCT |
| P434 | A_23_F | AAATCGATTGCCAACTGGACCAATACGATCAACTACT |
| P435 | A_23_R | CCAGTTGGCAATCGATTTGATGCCCAGGATCT |
| P438 | B_11_F_2 | TTAACGCGATGGGCAGCCAACCAGAAGAGCC |
| P439 | B_15_F_2 | TTAACGCGGTGGAAAGCCAACCAGAAGAGCC |
| P440 | B_22_F_2 | TTAACGCACGTGTGAGCCAACCAGAAGAGCC |
| P441 | B_22_R_2 | GGCTCACACGTGCGTTAAGGCTCCCCCAGGA |
| P442 | B_23_F_2 | TTAACTGGCGCCATAGCCAACCAGAAGAGCC |
| P443 | B_24_F_2 | TTAACGCGCGCTTTAGCCAACCAGAAGAGCC |
| P444 | B_26_F_2 | TTAACAGCGTGAGCAGCCAACCAGAAGAGCC |
| P445 | A_24_F_2 | GACAGCGCAGCCAACGGCACCAATACGATCAACTACT |
| P447 | A_24_R_2 | GCCGTTGGCTGCGCTGTCGATGCCCAGGATCT |
| P448 | A_25_F | GGCTCGAGCGCCAACTATACCAATACGATCAACTACT |
| P449 | A_25_R | ATAGTTGGCGCTCGAGCCGATGCCCAGGATCT |
| P450 | A_26_F | CTGTCGACCGCCAACAAAACCAATACGATCAACTACT |
| P451 | A_26_R | TTTGTTGGCGGTCGACAGGATGCCCAGGATCT |
| P452 | A_27_F | CGTTCGGCGGCCAACCATACCAATACGATCAACTACT |
| P453 | A_27_R | ATGGTTGGCCGCCGAACGGATGCCCAGGATCT |
| P454 | A_28_F | CATTCGATGGCCAACGAAACCAATACGATCAACTACT |
| P455 | A_28_R | TTCGTTGGCCATCGAATGGATGCCCAGGATCT |
| P456 | A_29_F | CAGTCGGTGGCCAACACCACCAATACGATCAACTACT |
| P457 | A_29_R | GGTGTTGGCCACCGACTGGATGCCCAGGATCT |
| P458 | A_30_F | AACTCGAAAGCCAACGATACCAATACGATCAACTACT |
| P459 | A_30_R | ATCGTTGGCTTTCGAGTTGATGCCCAGGATCT |
| P460 | A_31_F | GAATCGCATGCCAACAGCACCAATACGATCAACTACT |
| P461 | A_31_R | GCTGTTGGCATGCGATTCGATGCCCAGGATCT |
| P462 | A_32_F | AGCTCGCGTGCCAACTTTACCAATACGATCAACTACT |
| P463 | A_32_R | AAAGTTGGCACGCGAGCTGATGCCCAGGATCT |
| P464 | A_33_F | TGGTCGCGCGCCAACGAAACCAATACGATCAACTACT |
| P465 | A_33_R | TTCGTTGGCGCGCGACCAGATGCCCAGGATCT |
| P466 | A_34_F | GCGTCGCTGGCCAACTTTACCAATACGATCAACTACT |
| P467 | A_34_R | AAAGTTGGCCAGCGACGCGATGCCCAGGATCT |
| P468 | A_35_F | AGCTCGTGGGCCAACAGCACCAATACGATCAACTACT |
| P469 | A_35_R | GCTGTTGGCCCACGAGCTGATGCCCAGGATCT |
| P470 | A_36_F | ATTTCGGAAGCCAACAACACCAATACGATCAACTACT |
| P471 | A_36_R | GTTGTTGGCTTCCGAAATGATGCCCAGGATCT |
| P472 | A_37_F | ATTTCGCTGGCCAACCATACCAATACGATCAACTACT |
| P473 | A_37_R | ATGGTTGGCCAGCGAAATGATGCCCAGGATCT |
| P474 | A_38_F | CTGTCGATGGCCAACGATACCAATACGATCAACTACT |
| P475 | A_38_R | ATCGTTGGCCATCGACAGGATGCCCAGGATCT |
| P476 | A_39_F | ATTTCGGCGGCCAACAACACCAATACGATCAACTACT |
| P477 | A_39_R | GTTGTTGGCCGCCGAAATGATGCCCAGGATCT |
| P478 | A_40_F | CTGTCGATGGCCAACTTTACCAATACGATCAACTACT |

| | | |
|---|---|---|
| P479 | A_40_R | AAAGTTGGCCATCGACAGGATGCCCAGGATCT |
| P480 | A_41_F | ATTTCGTATGCCAACATTACCAATACGATCAACTACT |
| P481 | A_41_R | AATGTTGGCATACGAAATGATGCCCAGGATCT |
| P482 | A_42_F | AGCTCGACCGCCAACGAAACCAATACGATCAACTACT |
| P483 | A_42_R | TTCGTTGGCGGTCGAGCTGATGCCCAGGATCT |
| P484 | A_43_F | GATTCGGATGCCAACACCACCAATACGATCAACTACT |
| P485 | A_43_R | GGTGTTGGCATCCGAATCGATGCCCAGGATCT |
| P486 | A_44_F | GAATCGCAGGCCAACAAAACCAATACGATCAACTACT |
| P487 | A_44_R | TTTGTTGGCCTGCGATTCGATGCCCAGGATCT |
| P488 | A_45_F | AACTCGCAGGCCAACTATACCAATACGATCAACTACT |
| P489 | A_45_R | ATAGTTGGCCTGCGAGTTGATGCCCAGGATCT |
| P490 | A_46_F | AGCTCGGATGCCAACCGCACCAATACGATCAACTACT |
| P491 | A_46_R | GCGGTTGGCATCCGAGCTGATGCCCAGGATCT |
| P492 | A_47_F | CAGTCGCTGGCCAACGGCACCAATACGATCAACTACT |
| P493 | A_47_R | GCCGTTGGCCAGCGACTGGATGCCCAGGATCT |
| P494 | A_48_F | AAATCGGATGCCAACATTACCAATACGATCAACTACT |
| P495 | A_48_R | AATGTTGGCATCCGATTTGATGCCCAGGATCT |
| P501 | TB_1_F | TTAACGTGAGCGACAGCCAACCAGAAGAGCCTT |
| P502 | TB_1_R | GGCTGTCGCTCACGTTAAGGCTCCCCCAGGA |
| P503 | TB_2_F | TTAACTTCAGCTATAGCCAACCAGAAGAGCCTT |
| P504 | TB_2_R | GGCTATAGCTGAAGTTAAGGCTCCCCCAGGA |
| P505 | TB_3_F | TTAACTTCAGCACCAGCCAACCAGAAGAGCCTT |
| P506 | TB_3_R | GGCTGGTGCTGAAGTTAAGGCTCCCCCAGGA |
| P507 | TB_4_F | TTAACTTCACCACGAGCCAACCAGAAGAGCCTT |
| P508 | TB_4_R | GGCTCGTGGTGAAGTTAAGGCTCCCCCAGGA |
| P509 | TB_5_F | TTAACGTGAGCTACAGCCAACCAGAAGAGCCTT |
| P510 | TB_5_R | GGCTGTAGCTCACGTTAAGGCTCCCCCAGGA |
| P511 | TB_6_F | TTAACTTCCAGGAAAGCCAACCAGAAGAGCCTT |
| P512 | TB_6_R | GGCTTTCCTGGAAGTTAAGGCTCCCCCAGGA |
| P513 | TB_7_F | TTAACTTTACCTATAGCCAACCAGAAGAGCCTT |
| P514 | TB_7_R | GGCTATAGGTAAAGTTAAGGCTCCCCCAGGA |
| P515 | TB_8_F | TTAACATGAGCTATAGCCAACCAGAAGAGCCTT |
| P516 | TB_8_R | GGCTATAGCTCATGTTAAGGCTCCCCCAGGA |
| P517 | TB_9_F | TTAACTTCGCATATAGCCAACCAGAAGAGCCTT |
| P518 | TB_9_R | GGCTATATGCGAAGTTAAGGCTCCCCCAGGA |
| P519 | TB_10_F | TTAACGTTAGCACCAGCCAACCAGAAGAGCCTT |
| P520 | TB_10_R | GGCTGGTGCTAACGTTAAGGCTCCCCCAGGA |
| P521 | TB_11_F | TTAACTTCAGCCCGAGCCAACCAGAAGAGCCTT |
| P522 | TB_11_R | GGCTCGGGCTGAAGTTAAGGCTCCCCCAGGA |
| P523 | TB_12_F | TTAACGTGCAGACCAGCCAACCAGAAGAGCCTT |
| P524 | TB_12_R | GGCTGGTCTGCACGTTAAGGCTCCCCCAGGA |
| P525 | TB_13_F | TTAACATGCAGTATAGCCAACCAGAAGAGCCTT |
| P526 | TB_13_R | GGCTATACTGCATGTTAAGGCTCCCCCAGGA |
| P527 | TB_14_F | TTAACTTTAGCGAAAGCCAACCAGAAGAGCCTT |
| P528 | TB_14_R | GGCTTTCGCTAAAGTTAAGGCTCCCCCAGGA |
| P529 | TB_15_F | TTAACTTTGTGGAAAGCCAACCAGAAGAGCCTT |
| P530 | TB_15_R | GGCTTTCCACAAAGTTAAGGCTCCCCCAGGA |
| P531 | TB_16_F | TTAACTTTGCAACCAGCCAACCAGAAGAGCCTT |
| P532 | TB_16_R | GGCTGGTTGCAAAGTTAAGGCTCCCCCAGGA |
| P533 | TB_17_F | TTAACGTGAGCGAAAGCCAACCAGAAGAGCCTT |
| P534 | TB_17_R | GGCTTTCGCTCACGTTAAGGCTCCCCCAGGA |
| P535 | TB_18_F | TTAACATGCAGGAAAGCCAACCAGAAGAGCCTT |
| P536 | TB_18_R | GGCTTTCCTGCATGTTAAGGCTCCCCCAGGA |
| P537 | TB_19_F | TTAACATGAGCACGAGCCAACCAGAAGAGCCTT |
| P538 | TB_19_R | GGCTCGTGCTCATGTTAAGGCTCCCCCAGGA |
| P539 | TB_20_F | TTAACTTCGCAGAAAGCCAACCAGAAGAGCCTT |
| P540 | TB_20_R | GGCTTTCTGCGAAGTTAAGGCTCCCCCAGGA |
| P541 | TB_21_F | TTAACTTCCAGTATAGCCAACCAGAAGAGCCTT |
| P542 | TB_21_R | GGCTATACTGGAAGTTAAGGCTCCCCCAGGA |
| P543 | TB_22_F | TTAACGTGAGCTTTAGCCAACCAGAAGAGCCTT |
| P544 | TB_22_R | GGCTAAAGCTCACGTTAAGGCTCCCCCAGGA |
| P545 | TB_23_F | TTAACATGAGCGAAAGCCAACCAGAAGAGCCTT |
| P546 | TB_23_R | GGCTTTCGCTCATGTTAAGGCTCCCCCAGGA |
| P547 | TB_24_F | TTAACTTTAGCGATAGCCAACCAGAAGAGCCTT |
| P548 | TB_24_R | GGCTATCGCTAAAGTTAAGGCTCCCCCAGGA |
| P549 | TB_25_F | TTAACGTGCAGTATAGCCAACCAGAAGAGCCTT |
| P550 | TB_25_R | GGCTATACTGCACGTTAAGGCTCCCCCAGGA |
| P551 | TB_26_F | TTAACTTTAGCCATAGCCAACCAGAAGAGCCTT |
| P552 | TB_26_R | GGCTATGGCTAAAGTTAAGGCTCCCCCAGGA |
| P553 | TB_27_F | TTAACTTTAGCAACAGCCAACCAGAAGAGCCTT |

| | | |
|---|---|---|
| P554 | TB_27_R | GGCTGTTGCTAAAGTTAAGGCTCCCCCAGGA |
| P555 | TB_28_F | TTAACTTTAGCGGCAGCCAACCAGAAGAGCCTT |
| P556 | TB_28_R | GGCTGCCGCTAAAGTTAAGGCTCCCCCAGGA |
| P557 | TB_29_F | TTAACTTCAGCTTTAGCCAACCAGAAGAGCCTT |
| P558 | TB_29_R | GGCTAAAGCTGAAGTTAAGGCTCCCCCAGGA |
| P559 | TB_30_F | TTAACATGAGCCCGAGCCAACCAGAAGAGCCTT |
| P560 | TB_30_R | GGCTCGGGCTCATGTTAAGGCTCCCCCAGGA |
| P561 | TB_31_F | TTAACTTTCTGGAAAGCCAACCAGAAGAGCCTT |
| P562 | TB_31_R | GGCTTTCCAGAAAGTTAAGGCTCCCCCAGGA |
| P563 | TB_32_F | TTAACATGGTGGAAAGCCAACCAGAAGAGCCTT |
| P564 | TB_32_R | GGCTTTCCACCATGTTAAGGCTCCCCCAGGA |
| P565 | TA_1_F | CATTCGCTGGCCAACCAGACCAATACGATCAACTACT |
| P566 | TA_1_R | CTGGTTGGCCAGCGAATGGATGCCCAGGATCT |
| P567 | TA_2_F | CATTCGCTGGCCAACCCGACCAATACGATCAACTACT |
| P568 | TA_2_R | CGGGTTGGCCAGCGAATGGATGCCCAGGATCT |
| P569 | TA_3_F | CATTCGCTGGCCAACTGCACCAATACGATCAACTACT |
| P570 | TA_3_R | GCAGTTGGCCAGCGAATGGATGCCCAGGATCT |
| P571 | TA_4_F | CATTCGCCGGCCAACTTCACCAATACGATCAACTACT |
| P572 | TA_4_R | GAAGTTGGCCGGCGAATGGATGCCCAGGATCT |
| P573 | TA_5_F | CATTCGCTGGCCAACGATACCAATACGATCAACTACT |
| P574 | TA_5_R | ATCGTTGGCCAGCGAATGGATGCCCAGGATCT |
| P575 | TA_6_F | CAGTCGCTGGCCAACGTGACCAATACGATCAACTACT |
| P576 | TA_6_R | CACGTTGGCCAGCGACTGGATGCCCAGGATCT |
| P577 | TA_7_F | CATTCGCTGGCCAACTGGACCAATACGATCAACTACT |
| P578 | TA_7_R | CCAGTTGGCCAGCGAATGGATGCCCAGGATCT |
| P579 | TA_8_F | CATTCGCTGGCCAACCTGACCAATACGATCAACTACT |
| P580 | TA_8_R | CAGGTTGGCCAGCGAATGGATGCCCAGGATCT |
| P581 | TA_9_F | CATTCGCTGGCCAACTATACCAATACGATCAACTACT |
| P582 | TA_9_R | ATAGTTGGCCAGCGAATGGATGCCCAGGATCT |
| P583 | TA_10_F | CATTCGCTGGCCAACAAAACCAATACGATCAACTACT |
| P584 | TA_10_R | TTTGTTGGCCAGCGAATGGATGCCCAGGATCT |

Table S3. Oligonucleotides used in this study.
